# Sex-specific variation in the use of vertical habitat by a resident Antarctic top predator

**DOI:** 10.1101/2020.06.15.152009

**Authors:** Theoni Photopoulou, Karine Heerah, Jennifer Pohle, Lars Boehme

## Abstract

Patterns of habitat use are commonly studied in horizontal space, but this does not capture the four-dimensional nature of ocean habitats. There is strong seasonal variation in vertical ocean structuring, particularly at the poles, and deep-diving marine animals encounter a range of oceanographic conditions. We use hidden Markov models fitted to telemetry data from an air-breathing top predator to identify different diving behaviours and understand usage patterns of vertically distributed habitat. We show that preference for oceanographic conditions in the Weddell Sea, Antarctica, varies by sex in Weddell seals, and present the first evidence that both sexes use high-density, continental shelf water masses. Males spend more time in the colder, unique high-salinity shelf water masses found at depth, while females also venture off the continental shelf and visit warmer, shallower pelagic water masses. Both sexes exhibit a diurnal pattern in diving behaviour that persists from austral autumn into winter. These findings provide insights into the Weddell Sea shelf and open ocean ecosystem from a top predator perspective. The differences in habitat use in a resident, sexually monomorphic Antarctic top predator suggest a different set of needs and constraints operating at the intraspecific level, which are not driven by body size.

## 2 Background

Understanding what parts of an ecosystem are important for species is a cornerstone of ecological research. Important habitat is often detected by proxy; if a species regularly occurs in a habitat, it must fulfill a life-history function. For large marine vertebrates, occurrence is usually measured using location data from animal-attached instruments. However, identifying the drivers of marine population distributions from horizontal location data alone can be problematic for air-breathing deep-diving marine animals [1, 2]. This group spend most of their time underwater and are intrinsically difficult to observe. Depth is a fundamental dimension of their movement, and information is lost if dives are not considered. Vertical structuring of ocean habitats can enhance productivity and create predictable concentrations of resources (e.g., [3, 4, 5]). Deep-divers can benefit from this increased prey density at steep physical gradients [6, 7, 8, 9, 10, 11, 12, 13] and track its seasonality [14], which is especially strong at the poles (e.g., [15]). For most species of deep-diving wide-ranging marine vertebrates, we do not have a detailed understanding of what prey they consume nor the structure and functioning of the ecosystems that support them.

For air-breathing divers like pinnipeds, seabirds and cetaceans, dives are the result of the separation of two basic resources: air at the surface and prey resources at depth. Greater depths are more costly from a time-budget perspective, since transit likely excludes foraging [16], and physiologically more costly [17, 18, 19], due to the metabolic requirements of hunting and digestion. It follows that the habitat at these depth layers is expected to be profitable in terms of prey resources ([16], but also [20, 21]).

Understanding the environmental context of diving is key to linking behaviour to habitat and prey. Recent examples of integrative multi-species studies that describe the spatial distribution of different marine animal guilds include [22, 23, 24, 25]. However, surface variables do not reflect conditions at depth [26], and the methodology for relating diving behaviour to depth-varying environmental variables is underdeveloped in comparison. The collection of behavioural and *in situ* environmental data by satellite-linked animal-borne devices [27, 28] is a way of filling this data-gap and allows us to relate depth-varying behaviour to depth-varying conditions. In this study we characterise the relationship between diving behaviour and seawater density regimes (temperature and salinity) in a sexually monomorphic Antarctic top predator, the Weddell seal (*Leptonychotes weddellii*), in its namesake shelf sea. We examine sex-specific differences, as well as seasonal and diurnal trends, to understand intraspecific partitioning of resources. Although this is a single-species study, the approach can be used equally effectively for multi-species data. Incorporating diving animals’ use of depth is critical for making good predictions of future distributions (e.g., [29, 30]).

The southern Weddell Sea is a large embayment in the Atlantic sector of the Southern Ocean. It stands out among Antarctica’s shelf seas with a wide continental shelf and extensive winter sea ice production, which leads to the formation of high-salinity water. This creates a unique environmental context. Weddell Sea hydrography has been the focus of intense research to understand the mechanisms that underpin the formation of globally important dense bottom water [15, 31, 32, 33]. Weddell seals occur all around the Antarctic coast and their diving behaviour has been studied since the earliest years of animal-borne instrument development [34, 35]. Due to higher accessibility, most of what we know about Weddell seal foraging ecology comes from East Antarctica. In these areas, seals seldom venture off the continental shelf where a much warmer and a little fresher water mass - modified Circumpolar Deep Water - plays a central role [36, 37]. In contrast, the southern Weddell Sea is only accessible by ship during the austral summer and seals largely occur on sea ice. The interaction of Weddell seals with the hydrographically complex and varied Weddell Sea vertical habitat is not currently understood (but see [38, 39, 40]) and is likely to be different, owing to the availability of different water masses and deep ocean habitat. Previous work on foraging ecology in the Weddell Sea has found that dives target the surface, shallow epipelagic, and the mesopelagic region [38, 41, 42, 43], while commuting through intermediate depths. Although some pioneering work has been done to find out what occurs at these depths [38, 41, 42], knowledge of Weddell Sea shelf ecosystem remains poor.

The high degree of serial correlation in animal movement metrics makes it necessary to use analytical approaches that account for it. Hidden Markov models (HMMs) [44, 45] are time series models, which allow us to learn about the underlying process from multivariate observations (multiple data steams). They have been used to model diving data from several marine predators and are powerful for making inferences about complex temporal patterns and responses to environmental features [46, 47, 48, 49, 50]. We use HMMs to identify Weddell seal dive types and the effect of temporal covariates on the probability of switching between vertical movement states. Geographical differences in female and male diving activity [51] suggest that the two sexes may dive to different depths and use different water masses. We identify the depth at which seals spend time hunting and link hunting activity to *in situ* oceanographic conditions, making this the first study to examine sex differences in the relationship between at-depth environmental conditions and foraging in this circumpolar resident top predator in the southern Weddell Sea.

## 3 Methods

### 3.1 Data collection

Behaviour and oceanographic data were collected using telemetry instruments attached to the fur of adult Weddell seals in the southern Weddell Sea, Antarctica, after their annual molt. We used Conductivity-Temperature-Depth Satellite Relay Data Loggers (CTD-SRDLs) [52, 53], designed and manufactured by the Sea Mammal Research Unit Instrumentation Group (SMRU-IG), St Andrews, Scotland, UK. CTD-SRDLs employ the CLS Argos satellite system to relay data [54]. This produced a dataset from 19 Weddell seals (10 females, 9 males) instrumented in mid-February 2011 (Fig 1), as part of an oceanographic research program [15, 55]. This is the largest single-year telemetry dataset from this species in the Weddell Sea.

**Figure 1:**
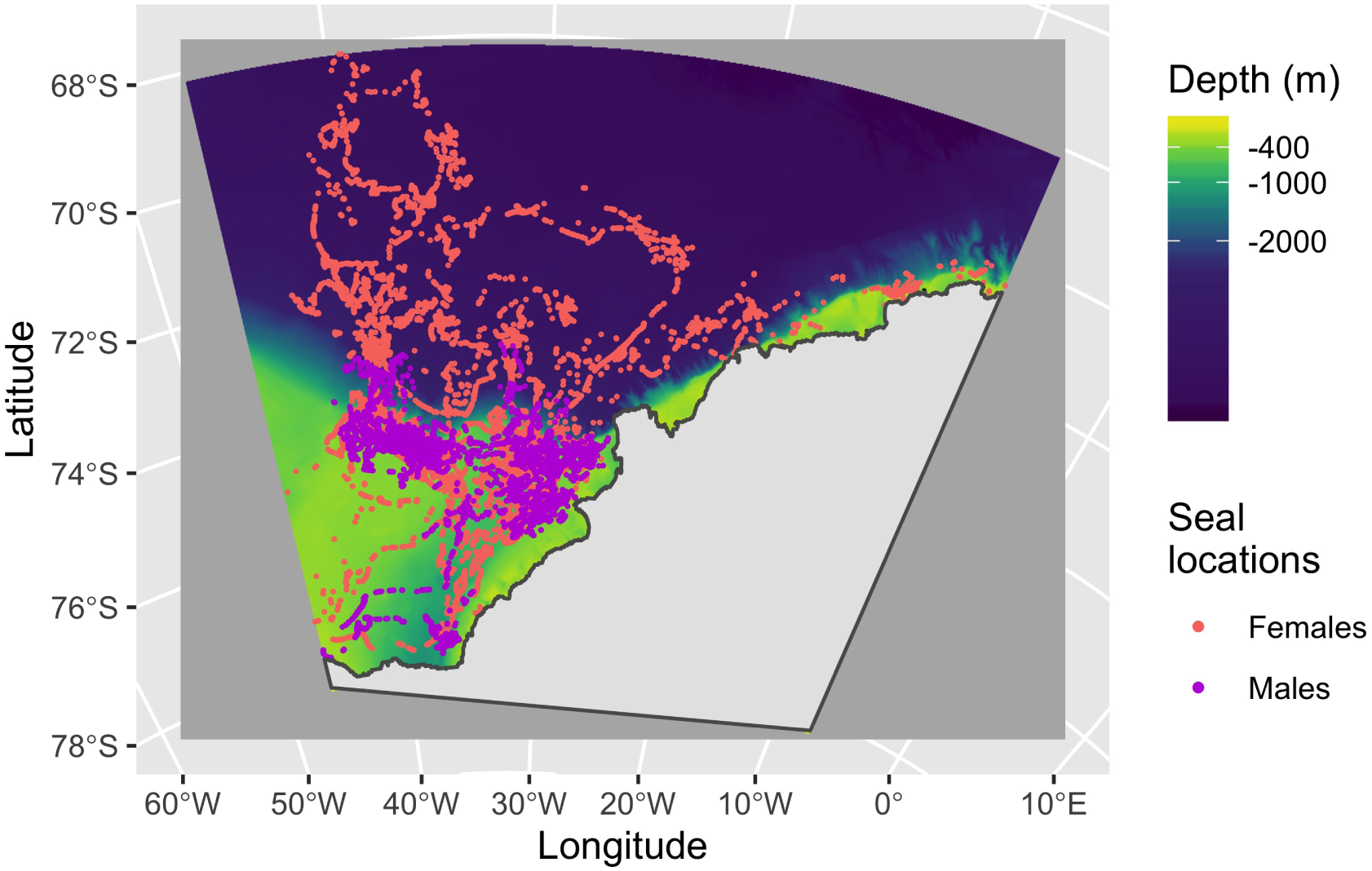
Satellite tracks from 19 instrumented Weddell seals carrying CTD-SRDL tags deployed in February 2011. The tracks are colour-coded by sex. The background colour represents the bathymetry (depth in metres at a 0.5km resolution). The colour bar has been scaled so that yellow areas represent the continental shelf and dark blue areas represent the deep ocean. There is a deep trough on the continental shelf at 40°W called the Filchner Depression. These are predicted locations from a correlated random walk model fitted to the original data, accounting for the estimated location error provided by CLS Argos, using the `foieGras` package in `R` [57].

A detailed description of the deployment technique and individual deployment data are presented in [56] and [51]. Dive data were collected using algorithms developed for high-latitude, deep-diving seals [52, 53, 58], and CTD data were collected, calibrated and processed according to the MEOP (Marine Mammals Exploring the Oceans Pole to Pole) consortium standards [27]. A summary of the data is presented in Supplementary Material S1.

### 3.2 Data processing

We consider a near-complete sequence of behaviour by combining the haulout, surface and dive information returned by the tag (for definitions see Supplementary Material S1). We do not consider short surface intervals between these events. We found that the amount of data reduces substantially after week 24 (13-19 June 2011) so we only consider data up to 19 June.

We calculated hunting depth and time spent hunting during dives using methods developed by [21]. Heerah *et al*. [21] used high-resolution time-depth data and triaxial accelerometer data from Weddell seals to detect prey capture attempts (PrCAs) and found a strong correlation between vertical sinuosity, swimming speed and the number of prey capture attempts. We use their estimated vertical speed threshold (0.5m/sec) to extract segments of dives with low vertical speeds (hunting segments) and calculate the total duration of time spent in “hunting” mode in each dive ([10] and Supplementary Material S2). We use the depth of the longest hunting segment, where most PrCAs occur, as the hunting depth in our analyses (Supplementary Material S2).

There is a small number of dives that have no hunting according to the vertical speed threshold (508 out of 20662 dives, 2.5%). In these cases the hunting-specific variables (e.g., depth and proportion of dive time spent hunting) have missing values.

Only some dive profiles have a simultaneous CTD profile. To overcome this, we interpolated variables linearly in time, between time points where CTD data were available, and assigned them to the intervening dive times (Supplementary Material S3.1). We extracted bathymetry at the location of each seal dive after correcting the tracks using a model-based approach (Supplementary Material S3.2).

### 3.3 Statistical analysis

We are primarily interested in describing diving behaviour, which includes foraging, so we consider the underlying movement state as known for haulouts and surface events. We allow additional states to describe different diving behaviours. It was obvious from exploratory data analysis that female and male seals display different movement modes, which are differently affected by covariates, and therefore warrant separate models. We use behavioural and environmental variables from each behavioural record to derive five data streams (state-dependent variables) to be modelled using a multivariate HMM. The state-dependent variables are 1) behaviour duration (sec), 2) hunting depth (m), 3) proportion of dive duration spent hunting (hunting time/duration: 0-1), 4) proportion of bathymetry reached at hunting depth (hunting depth/bathymetry: 0-1) (see Supplementary Material S3.3), and 5) salinity (psu) at hunting depth. Only behaviour duration is meaningful for non-dive behaviours, so this is the only state-dependent variable that contributes to the estimation of haulout and surface events. All state-dependent variables contribute to the estimation of dive states.

We fit two multivariate HMMs (females, males) to estimate underlying states in the time series of the five state-dependent variables, which are assumed to be contemporaneously conditionally independent given the states. We model behaviour duration and hunting depth using a gamma distribution for non-zero positive values, proportion of dive time spent hunting as a beta distribution for values between zero and one, proportion of bathymetry reached as a single probability between zero and one, and salinity as a normal distribution. Salinity values are strictly positive but far from zero, so parameters do not need to be bounded.

Based on exploratory data analysis and existing knowledge regarding Weddell seal diving behaviour [34, 59, 36, 40], we choose two covariates that act on the probability of transition between states. These include local time of day as a circular variable with two components, cosine and sine, and week of the year, as well as an interaction term between time of day and week. The model formulation (the state-dependent variables, the temporal covariates, and the way the covariates were included in the model) was chosen carefully, based on biological system knowledge, so we did not carry out covariate selection and the full model was used for both females and males (Supplementary Material S4).

We include the time series of salinity in the state process rather than as a covariate on the transition probabilities. Our rationale is that we regard the hydrographic conditions encountered during a dive of a particular type, to be part of that movement behaviour, not an external factor influencing it. Weddell seals are long-lived, slowly reproducing large mammals with complex spatial memories. It seems unlikely that they respond mechanically to the abiotic conditions they encounter from one dive to the next. We initially included temperature as a covariate but removed it because we found a complete overlap of the state-dependent distributions, suggesting it is not useful for separating out the groupings in the data. We estimate the parameters of the state-dependent distributions, the covariate effects, and transition probabilities using numerical maximisation of the likelihood, implemented in R [60], using the nlm function. The computation of the covariate-dependent transition probability matrices and the forward algorithm were coded in C++ (Supplementary Material S4).

We chose initial values empirically for each sex separately by looking at the distributions of the observed data, and used twenty sets of starting values to ensure the algorithm found a global maximum. We use the Viterbi algorithm to calculate the most likely state sequence for each individual.

## 4 Results

### 4.1 State-dependent distributions

The data from females and males were best described using different numbers of dive states. This resulted in a 5-state model for males and a 6-state model for females (Supplementary Material S4). In each case, the two non-diving states were provided as known states and dive states were estimated from the data (females: 4 dive states, males: 3).

The estimated state-dependent distributions displayed in Fig 2 show that female and male Weddell seals have three very similar vertical movement states. These are most easily thought of in terms of their hunting depth and proportion of bathymetry reached: 1) very short shallow dives, 2) slightly deeper dives in the epipelagic layer of the water column, and 3) deep dives with high probability of reaching the bottom, where they access high salinity water masses found only on the continental shelf (Table 1, Fig 2). Females also carry out a fourth dive type, in the pelagic layer of the water column. These dives are shorter than benthic dives and take place over the deep ocean, where the seals access lower salinity water masses (Table 1, Fig 2). In males, the pelagic dive state appears to be absent, although the epipelagic state is a bit deeper than in females. There was no compelling evidence of differences in proportion of time spent hunting between dive types or sexes.

**Table 1:** Parameter estimates for state-dependent distributions for female and male Weddell seals, from the respective HMMs - 6 states for females and 5 states for males. The proportion state occupancy is calculated as a the proportion of observations labelled as each state. Note that there are 4 dive states for females to spread their activity over compared to 3 in males. This naturally leads to smaller proportions in the 6-state scenario. However, if all states occur equally often, the proportions would all be smaller, which we do not see.

|  | Haulout events | Surface events | Shallow dives | Epipelagic dives | Pelagic dives | Benthic dives | Sex |
| --- | --- | --- | --- | --- | --- | --- | --- |
| Duration | 2.8hrs (2.7-3.0) | 37.9min (36.4-39.5) | 1.0min (59-64sec) | 7.4min (7.2-7.7) | 14.1min (13.9-14.2) | 19.0min (18.7-19.2) | F |
|  | 3.5hrs (3.3-3.7) | 34.3min (32.6-35.9) | 2.3min (2.2-2.4) | 11.0min (10.7-11.3) |  | 18.2min (18.1-18.4) | M |
| Depth (m) |  |  | 8.8 (8.6-8.9) | 38.5 (37.2-39.8) | 193.8 (189.0-198.6) | 413.1 (409.9-416.3) | F |
|  |  |  | 10.2 (10.1-10.4) | 90.0 (86.8-93.2) |  | 411.8 (408.5-415.1) | M |
| Proportion of dive spent hunting |  |  | 0.51 (0.50-0.51) | 0.55 (0.54-0.56) | 0.49 (0.48-0.50) | 0.43 (0.42-0.44) | F |
|  |  |  | 0.53 (0.52-0.53) | 0.46 (0.46-0.47) |  | 0.42 (0.41-0.42) | M |
| Proportion of bathymetry reached |  |  | 0.00 | 0.00 | 0.00 | 0.89 (0.72-1.00) | F |
|  |  |  | 0.00 | 0.00 |  | 0.91 (0.77-1.00) | M |
| Salinity at depth (psu) |  |  | 34.18 (34.178-34.192) | 34.29 (34.283-34.292) | 34.44 (34.433-34.444) | 34.61 (34.603-34.608) | F |
|  |  |  | 34.23 (34.226-34.234) | 34.36 (34.351-34.360) |  | 34.59 (34.591-34.598) | M |
| State occupancy | 0.13 (0.10-0.17) | 0.21 (0.18-0.28) | 0.17 (0.12-0.23) | 0.24 (0.14-0.36) | 0.15 (0.07-0.30) | 0.09 (0.01-0.26) | F |
|  | 0.13 (0.10-0.17) | 0.20 (0.12-0.26) | 0.31 (0.09-0.42) | 0.19 (0.12-0.31) |  | 0.17 (0.11-0.36) | M |

**Figure 2:**
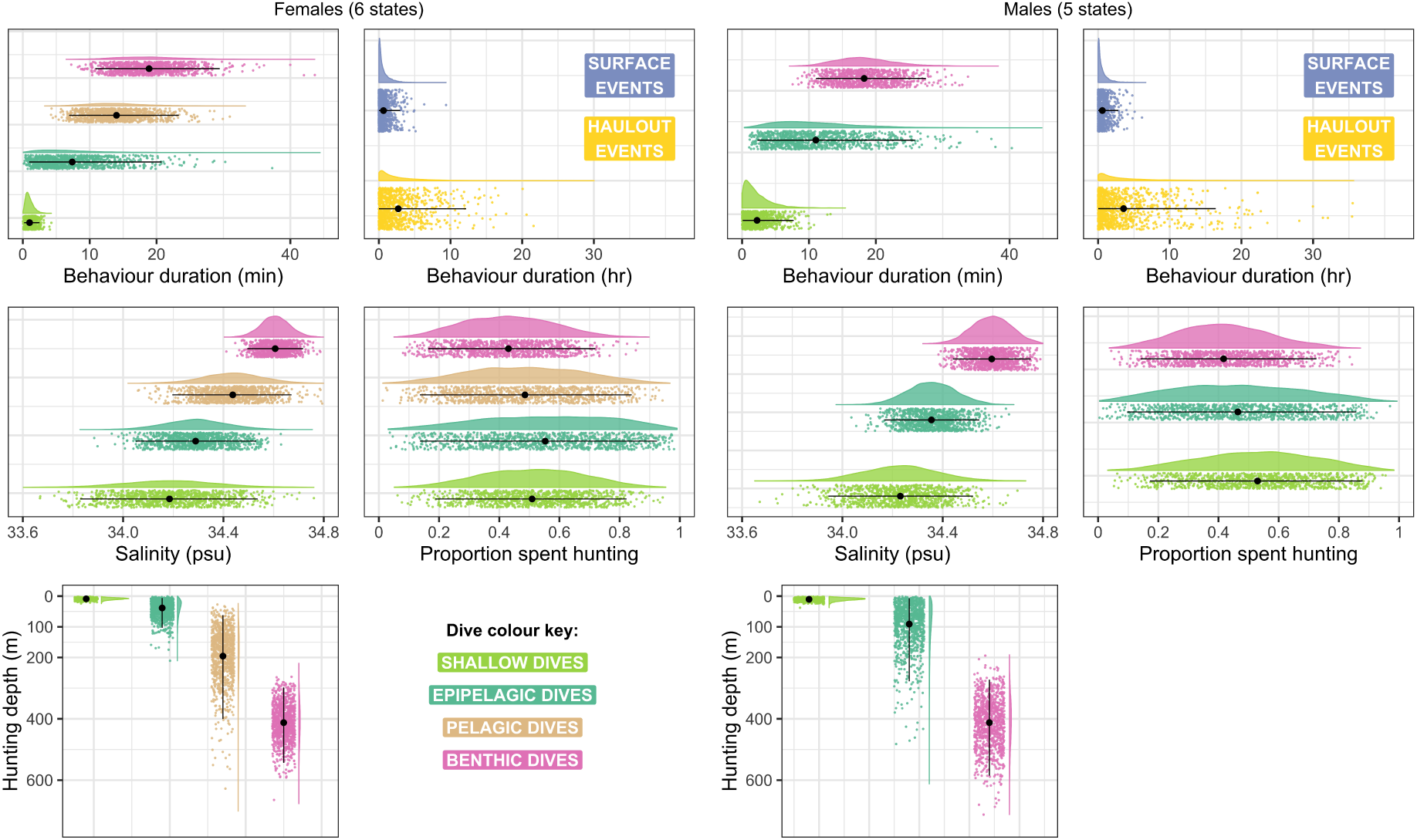
State density distributions for dive states (shallow, epipelagic, pelagic and benthic dives) and two non-dive states (haulout, surface) in female (left) and male (right) Weddell seals. The density distributions are colour-coded by state, with a cloud of points below them. The cloud represents a realisation of the estimated state-dependent distribution, based on a sample of 1000 points. The black point and line going through each coloured cloud of points represent the mean estimated by the model and the interval covering 95% of the probability mass. In females all 4 dive states are present, while in males there are 3 dive states (shallow, epipelagic and benthic).

Female and male deep dives are both benthic (Fig 2), with a 90% chance of reaching the bottom. This suggests that during these dives seals are targeting prey that occur near or at the bottom. The proportion of Viterbi decoded states differed between females and males (Table 1). Both sexes are equally likely to haul out and spend time at the surface, but males are almost twice as likely to carry out shallow dives (F: 17%, M: 31%) and benthic dives (F: 9%, M: 17%), while females are more likely to carry out epipelagic dives (F: 24%, M: 19%). Pelagic dives clearly make up a regularly used movement mode for females with 15% of records being estimated to belong to this state.

We checked the model fit by 1) computing the pseudo-residuals for each of the state-dependent variables, and by 2) simulating data from the model and checking how well it matched the observed data (Supplementary Material S5). There was no evidence of systematic lack of fit for either model.

### 4.2 Covariate effects

The distribution of state occupancy (i.e., the equilibrium probabilities as functions of covariates) is shown in Fig 3 (females top, males bottom). These plots represent the probability (y-axis) of finding a seal in a given state (columns), during a week of the year (rows) throughout the 24-hour cycle (x-axis). The grey ribbon represents the area enclosed by the lower and upper confidence bounds of the 95% interval of the estimated effect. We provide the state transition probabilities for the conditions shown in Figure 3, in Supplementary Material S5.

**Figure 3:**
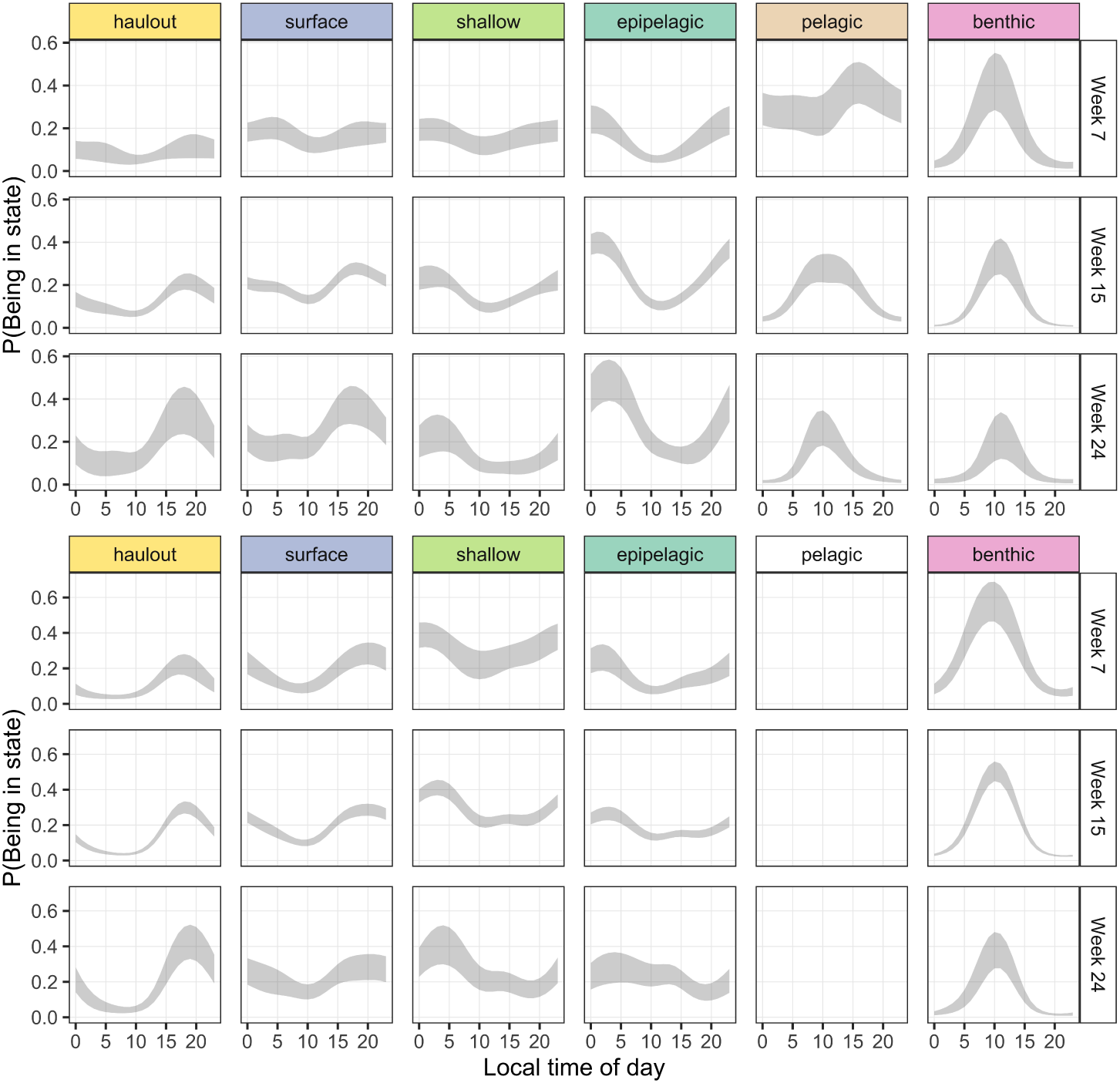
The probability (y-axis) of finding a seal in a given state (columns), during a week of the year (rows) throughout the 24-hour cycle (x-axis). The grey ribbon represents the area enclosed by the lower and upper confidence bounds of the 95% interval of the estimated effect, which were calculated using the delta method. Time of day corresponds to local time calculated from the location of each data point.

A seasonal effect is evident in female diving behaviour, particularly pelagic diving and benthic diving. Early in the year (week 7, late summer) pelagic dives are the most common movement mode, and occur throughout the day. Benthic dives are also common, but occur mainly in the day. By midwinter (week 24) deep dives are less common, replaced by nighttime epipelagic diving.

Male diving behaviour appears less seasonal. Benthic dives are dominant throughout the observation period and are most likely to occur around midday. Near midwinter, they become less dominant compared with other dive states, especially shallow dives.

The difference in oceanographic regimes used by female and male seals is clear in the temperature-salinity diagrams associated with their dive profiles. Although temperature was not included in the model we can still visualise the Viterbi decoded states in temperature-salinity (TS) space (Fig 4). The distribution of states in TS space is unsurprising for the shallower states (shallow, epipelagic dives) because there is a limited range of conditions that are found in these surface layers. However, the deeper states show a trend: females use Modified Warm Deep Water and Warm Deep Water extensively for pelagic dives. These water masses are found along the diagonal line seen in the female TS diagram. In contrast, males in this dataset almost never visited areas with at-depth water temperatures warmer than 0°C. The TS conditions associated with benthic dives are relatively consistent between females and males. In both plots, Ice Shelf Water and High Salinity Shelf Water, as well as its interface with precursors to Modified Warm Deep Water, are visited regularly (see Figure 3 in [31] for water mass definitions).

**Figure 4:**
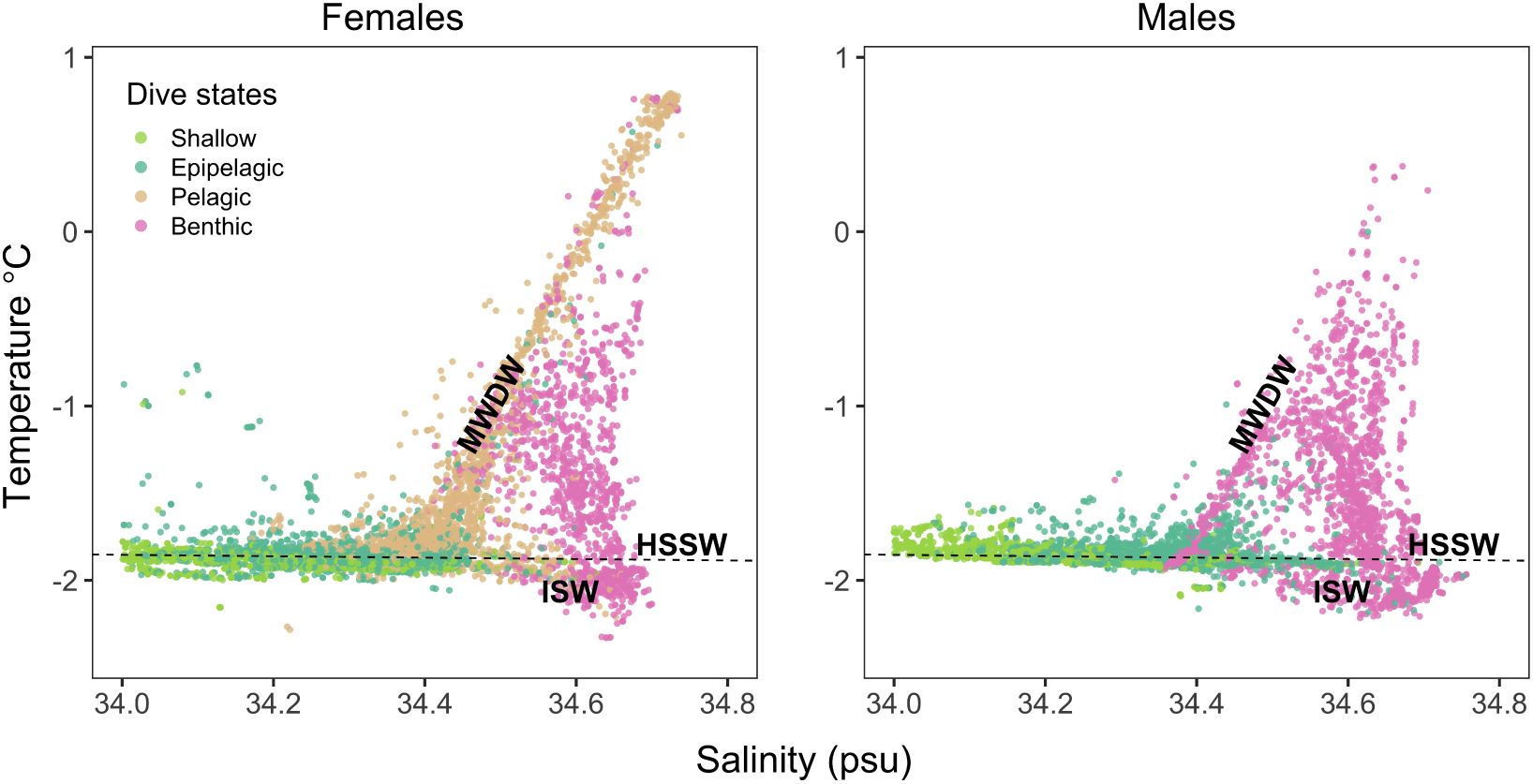
Temperature-salinity diagrams for dives from female and male seals coloured by Viterbi decoded state. The points in each plot correspond to the conditions encountered by seals at the hunting depth of each dive. The water mass labels correspond to High Salinity Shelf Water (HSSW), Ice Shelf Water (ISW) and Modified Warm Deep Water (MWDW). These labels are transcribed from Figure 3 of Nicholls *et al*. [31].

## 5 Discussion

The aim of this study is to characterise diving behaviour and vertical habitats used by Weddell seals in the southern Weddell Sea. We do this by modelling the relationship between diving and seasonal environmental conditions in an integrative way, using HMMs. We find two movement strategies, that in our dataset correspond to sex-specific differences. Females and males have some overlap in diving behaviour, but there are also important differences. Males mainly carry out benthic deep dives, consistent with their spatial location on the continental shelf into winter (Fig 1). In contrast, female deep dives are both benthic and pelagic, consistent with their extensive off-shelf autumn and winter distribution. In both sexes, deep dives happen during the day, centered on local noon, (Fig 3). This pattern is present throughout the seasons represented in our dataset, even when there is little light at local noon. Both sexes use high-salinity shelf water masses throughout, from February to June. Regular visits to this unique and inaccessible vertical habitat - the coldest possible liquid water habitat at −2°C - is worth the effort of carrying out deep dives, which have been found to be energetically costly [19]. Female diving behaviour follows a mixed seasonal pattern. In late summer, pelagic dives occur throughout the day, with a peak in the afternoon, while benthic dives are centered on midday. Near midwinter the dominant dive type is epipelagic diving. This is in contrast with the comparatively aseasonal diving behaviour of males.

Dive types represent different vertical habitat types. We found that male seals avoid the pelagic zone almost entirely. What does this suggest about the pelagic ecosystem on the southern Weddell Sea continental shelf? Dives down to the continental shelf benthos place the seal in some of the coldest, densest water in the world. Harcourt *et al*. [61] found that males that dived deeply during the breeding season lost weight slower than those that dived more shallowly. This is evidence that the prey resources accessed at depth are likely to be particularly profitable, hinting at the productivity of on-shelf Antarctic habitats [62]. The question then is, why do female seals leave the shelf and venture north? Perhaps due to over-crowding and density-dependent prey depletion, which females avoid by leaving the area. Another possible explanation is that the resources found off the continental shelf are as profitable, or even more so. Females can move north to exploit them at shallower, less energetically costly depths, unhindered by the need to establish territories for the autumn. This pattern is similar to southern elephant seals from Isles Kerguelen, where the males commonly stay in the sea ice and forage on the Antarctic continental shelf while females move north as winter progresses and forage in the marginal ice zone [63]. The life history constraints that likely bring about this pattern for sexually dimorphic elephant seals are reversed for sexually monomorphic Weddell seals - the males are limited by the need to establish territories, keeping them in the south - not the females. In both cases, the marginal ice zone is clearly seasonally attractive for top predators.

The seasonality in female movement behaviour manifests both latitudinally and vertically. In late summer, females are most likely to be found diving pelagically throughout the day, and benthically at midday. This is in contrast to what has been found in East Antarctica (see [59] but also [36]). The hydrographic conditions associated with pelagic dives are compellingly clear: Modified Warm Deep Water, mixing with early Winter Water, and the continuum up the temperature gradient towards Warm Deep Water (Fig 4 and Fig 3 in Nicholls *et al*. [31] for reference). These conditions have consistently been shown to be profitable to female phocid seals (e.g., [7, 36, 30, 11, 26, 40]). What pelagic resources are females exploiting within the Weddell Sea Gyre? Sea ice has a high concentration of detritus and living organisms, which contribute to year-round biological productivity and community development, particularly under older ice floes [64]. This leads to a cascade of productivity, and may give rise to the prey base exploited during the two shallow dive types we found. The existence of multi-year sea ice in the Weddell Sea may be the critical factor that allows female Weddell seals to forage successfully (and haul out) away from the coast over deep water. The post-moult period finds female Weddell seals in need of replenishing their energy reserves after a period of reduced foraging due to lactation and the annual moult. A shallower pelagic diving strategy could afford the added energetic benefit of minimizing dive costs.

The lack of seasonality in male diving behaviour raises questions about the prey base they are exploiting. Hydrographically, the end of summer sees the beginning of a reduction in the inflow of warm water onto the continental shelf in the southern Weddell Sea [55]. Warm water enters the shelf as Modified Warm Deep Water in the region of the Filchner Depression, especially the eastern and western edges of the Filchner Sill (Fig 6 and 7 in Nicholls *et al*. [31] and Årthun *et al*. [55]). These physical changes must have some ecosystem effects but are not so great as to induce a discernible change in diving behaviour. However, a reduction in the probability of benthic dives and increase in epipelagic dives hint towards a shift in reliance towards surface layers in winter. Changes not detectable by our approach include benthic prey type or prey behaviour. The TS diagram for males shows that some benthic dives occur in Modified Warm Deep Water (diagonal line starting at 34.4psu in Fig 4) but the clear majority of them occur in the denser, saltier water characteristic of sub-ice-shelf circulation and melting [65].

The shelf ecosystem that exists under these extreme temperature conditions is not well-studied, but is likely to experience a high degree of stability due to the narrow range of temperature and salinity conditions that can exist there [65]. It is known to support a large amount of biomass through Weddell seals and their likely prey, Antarctic toothfish (*Dissostichus mawsoni*) and other notothen fish (*Pleurogramma antarcticum, Trematomus loennbergii*). Stability and high biomass are attributes of mature ecosystems where large body size, narrow niche specialisation and long, complex life cycles are observed [66]. These attributes fit with what we know about the Weddell Sea continental shelf from a top predator perspective. Riotte-Lambert and Matthiopoulos [67] show that learning and memory are emergent properties of animals in a system with moderate-to-high environmental predictability. At slightly lower predictability, more social information use is favoured. Although the degree of social information exchange is not known for Weddell seals, they have a substantial underwater vocal repertoire known to function socially, at least in the breeding season. This would place the predictability of their environment on the mid-to-upper end of the scale (Fig 1 in [67]) at a time scale comparable with the lifetime of a Weddell seal. As non-migratory top predators, Weddell seals are likely to be instrumental in the stability of the Antarctic shelf ecosystem as a whole, forming a feedback loop between movement ecology and environmental predictability [67].

Historically, bottom trawls found high biomass at 600m on the southern slopes of the Filchner Depression was from notothen species (especially *D. mawsoni, P. antarcticum, T. loennbergii*) [62]. This is in contrast with the narrow eastern shelf where overall fish biomass was lower (Fig 3 in Ekau [62]), and which seals tend to avoid in our dataset. Over the Filchner Depression Antarctic toothfish were only caught in summer at 420-670m depth and dominated in terms of biomass, along with *T. loennbergii* and large specimens of Antarctic silverfish (*P. antarcticum*). The distribution and temporal availability of these high-biomass, large-bodied fish is highly consistent with the location and dive depth of Weddell seals on the shelf. In other parts of their range, Weddell seals have been found to mainly consume *P. antarcticum*, with seasonally varying contributions of larger prey such as *D. mawsoni* [68]. Goetz *et al*. [68] suggest that the proportional contribution of larger prey may increase with age. We do not have age information for the individuals tagged in this study, but it is possible that there is further intraspecific niche separation beyond sexual segregation.

A diurnal dive pattern was observed early on in Ross Sea Weddell seals [34] and more recently in East Antarctica [36] and the Weddell Sea [40], with seals diving deeply in the day and shallowly at night. The shallow depths visited during nighttime are now known to overlap with the diel vertical migration of *P. antarcticum* [69, 36]. The depths and dive types described in [34, 70] correspond closely to the four dive states we identify for females. In agreement with previous studies, we find that epipelagic (*<*100m) dives are more likely at night [34, 71, 36, 40]. The daytime pelagic (*<*250m) female dives deviate from what we expected for pelagic diving behaviour based on other species (e.g., southern elephant seals [7]). The pelagic ecosystem in the middle of the Weddell Sea Gyre is even less studied than the shelf benthos, but it seems likely that at these depths (*∼* 200m) seals are foraging on some age class of *P. antarcticum* and other notothen species in the day, and moving shallower at night (*∼* 50m).

The biological inferences we have drawn hinge on our analytical approach. The major advantages of HMMs for diving are that with them we can combine many aspects of diving behaviour and make inferences about possible behavioural classes in multiple dimensions, e.g., depth, duration, hunting behaviour, distance from features, physiological state, etc. A drawback of HMMs based on data collected in the wild, without direct observations, is that we do not know how well our state interpretations correspond to reality. If we repeated the study, increasing the sample size, measuring physiological parameters relating to life stage or energetic status, relatedness, age and, of course, prey field, would help shed light on some of the unanswered questions. The need for information on prey field, prey type and prey profitability is also raised by the lack of signal in the proportion of time spent hunting.

## 6 Conclusion

In this study we show that Weddell Sea Weddell seals segregate in horizontal, vertical and hydrographic space from late summer into midwinter. The residency of males on the continental shelf, and the lack of pelagic diving, suggest that benthic and shallow under-ice habitats provide adequate prey resources to support this top predator. Whether by competition avoidance, or in search of “better” (more reliable, abundant, energetically profitable?) foraging opportunities, most of the females in our dataset left the shelf and moved north over the abyssal plain, supported by pelagic and epipelagic resources, before returning east and south towards shelf areas. These movements are associated with access to different hydrographic conditions, which are exploited via the diving patterns we have documented here. This is the earliest evidence both of 1) sex-specific diving patterns in this monomorphic top predator, and of 2) their common reliance on the high-salinity shelf water masses unique to the southern Weddell Sea.

## 7 Acknowledgements

We would like to thank Marius Årthun, and the officers and crew of RRS Ernest Shackleton for their support during cruise ES054. Special thanks to Keith Nicholls for valuable input on Weddell Sea oceanography, and Mike Fedak for productive discussions on pinniped foraging strategies. The data collection was funded by NERC grants NE/G014833/1 and NE/G014086/1. TP was supported by a Royal Society Newton International Fellowship (NF170682). KH was supported by a Marie-Sklodowska Curie Research Fellowship. The oceanographic data were processed and made freely available by the International MEOP Consortium and the national programs that contribute to it. (http://www.meop.net).

## 8 Author contributions

TP and LB collected the data and conceived the study. TP, JP and KH developed the methods. TP carried out the analysis and wrote the manuscript. KH carried out the analysis in Supplementary Material S2. JP provided high level statistical support. All authors contributed critically to the manuscript and gave approval for publication.

## 9 Ethics and permits

Animal capture and handling protocols were reviewed and approved by the University of St Andrews Teaching and Research Ethics Committee (UTREC) and the Animal Welfare and Ethics Committee (AWEC) as part of the ethical review process. Permission to work in the Antarctic was granted under permit No.S7-4/2010 of the Antarctic Act 1994. Capture and deployment of satellite transmitters was carried out by experienced personnel with UK Animal (Scientific Procedures) Act 1986 Personal Licenses.

## 10 Availability of data and material

The dataset analysed for this article is available in a Zenodo repository at http://doi.org/10.5281/zenodo.3820359 and R code for running the models can be found on GitHub at https://github.com/theoniphotopoulou/emews.

