## Supplementary material 1 for "Sex-specific variation in the use of vertical habitat by a resident Antarctic top predator"

| Seal ID | Sex | Start date | End date | Dive duration (min) | Maximum dive depth (m) | Hunting depth (m) | Proportion of dive time spent hunting | $\frac{Huntingdepth}{Bathymetry}$ | Dives without hunting |
|---|---|---|---|---|---|---|---|---|---|
| ct70-356-11 | M | 08 Feb | 22 Feb | 15.5±7.1 (0.5-27.0) | 331±178 (5-574) | 295±186 (5-574) | 0.46±0.17 (0.00-0.81) | 0.57±0.36 (0-1.00) | 4/168 |
| ct70-486-11 | M | 09 Feb | 03 Aug | 6.6±6.9 (0.5-29.0) | 112±158 (5-594) | 99±148 (5-569) | 0.51±0.19 (0.00-0.94) | 0.23±0.36 (0-1.00) | 18/1038 |
| ct70-488-11 | F | 10 Feb | 07 Oct | 12.5±9.3 (0.5-50.3) | 187±190 (5-844) | 171±187 (5-724) | 0.54±0.18 (0.00-0.97) | 0.39±0.42 (0-1.00) | 14/1700 |
| ct70-490-11 | M | 09 Feb | 01 Oct | 6.1±7.0 (0.5-28.5) | 101±160 (5-694) | 93±157 (5-689) | 0.50±0.20 (0.00-0.94) | 0.19±0.35 (0-1.00) | 15/839 |
| ct70-491-11 | M | 15 Feb | 13 Oct | 11.1±8.3 (0.5-47.3) | 161±219 (5-874) | 128±203 (5-809) | 0.44±0.24 (0.00-0.94) | 0.25±0.35 (0-1.00) | 59/1683 |
| ct70-499-11 | M | 11 Feb | 10 Nov | 7.7±7.4 (0.5-37.3) | 140±170 (5-724) | 122±160 (5-639) | 0.46±0.19 (0.00-0.94) | 0.29±0.38 (0-1.00) | 35/1367 |
| ct70-500-11 | M | 11 Feb | 03 Oct | 8.1±7.2 (0.5-48.3) | 162±175 (5-574) | 149±169 (5-549) | 0.46±0.18 (0.00-0.94) | 0.35±0.40 (0-1.00) | 24/1033 |
| ct70-501-11 | F | 08 Feb | 10 Oct | 9.2±6.7 (0.5-39.3) | 121±119 (5-584) | 101±107 (5-584) | 0.55±0.19 (0.00-0.97) | 0.04±0.07 (0-1.00) | 24/1555 |
| ct70-502-11 | F | 09 Feb | 27 Mar | 9.0±7.1 (0.5-40.3) | 192±183 (5-594) | 177±178 (5-594) | 0.45±0.18 (0.00-0.97) | 0.41±0.42 (0-1.00) | 14/506 |
| ct70-503-11 | F | 08 Feb | 19 Sep | 7.7±5.3 (0.5-34.3) | 129±115 (5-614) | 110±104 (5-614) | 0.45±0.21 (0.00-0.90) | 0.09±0.16 (0-1.00) | 49/1175 |
| ct70-526-11 | M | 10 Feb | 01 Jun | 8.7±7.7 (0.5-34.3) | 152±165 (5-564) | 140±164 (5-519) | 0.48±0.19 (0.00-0.97) | 0.35±0.41 (0-1.00) | 26/1211 |
| ct70-633-11 | M | 14 Feb | 19 Sep | 8.3±6.7 (0.5-28.0) | 149±170 (5-654) | 135±166 (5-654) | 0.45±0.21 (0.00-0.87) | 0.25±0.33 (0-1.00) | 34/871 |
| ct70-634-11 | M | 10 Feb | 12 Jul | 10.6±8.7 (0.5-45.3) | 188±182 (5-1269) | 173±178 (5-569) | 0.50±0.17 (0.00-0.90) | 0.42±0.43 (0-1.00) | 24/1364 |
| ct70-637-11 | F | 14 Feb | 08 Oct | 7.8±5.7 (0.5-34.3) | 119±133 (5-844) | 98±119 (5-519) | 0.49±0.21 (0.00-0.94) | 0.10±0.15 (0-0.98) | 24/879 |
| ct70-638-11 | F | 09 Feb | 24 Aug | 5.2±4.5 (0.5-23.5) | 83±109 (5-461) | 72±101 (5-421) | 0.50±0.22 (0.00-0.94) | 0.10±0.23 (0-1.00) | 22/1018 |
| ct70-640-11 | F | 13 Feb | 29 Oct | 9.4±9.2 (0.5-95.3) | 122±153 (5-614) | 102±140 (5-609) | 0.51±0.21 (0.00-0.97) | 0.20±0.30 (0-1.00) | 28/1356 |
| ct70-642-11 | F | 11 Feb | 04 May | 7.0±5.4 (0.5-25.0) | 147±162 (5-704) | 127±156 (5-699) | 0.42±0.22 (0.00-0.93) | 0.26±0.33 (0-1.00) | 46/910 |
| ct70-643-11 | F | 12 Feb | 01 Nov | 7.1±5.8 (0.5-28.5) | 85±106 (5-594) | 70±96 (5-564) | 0.52±0.20 (0.00-0.90) | 0.03±0.08 (0-1.00) | 26/1308 |
| ct70-650-11 | F | 10 Feb | 28 Jul | 10.8±8.7 (0.5-38.3) | 184±194 (5-734) | 165±188 (5-729) | 0.50±0.19 (0.00-0.97) | 0.35±0.40 (0-1.00) | 22/1190 |

Table S1. Descriptive statistics regarding dive parameters from the individual Weddell seals used in the main analysis. All seals were instrumented in the southern Weddell Sea and all dates refer to the year 2011. Dive parameters are presented as mean ± 1 standard deviation and the range is given in brackets in grey. Some dives did not have any segments that were identified as hunting segments using the methods developed by [1], these number of these dives is shown in the last column as a fraction of the total number of dives recorded for each animal. When no hunting segments were recorded there are missing values for hunting depth, proportion of dive spent hunitng, environmental variables (temperature and salinity) and the proportion of bathymetry reached.

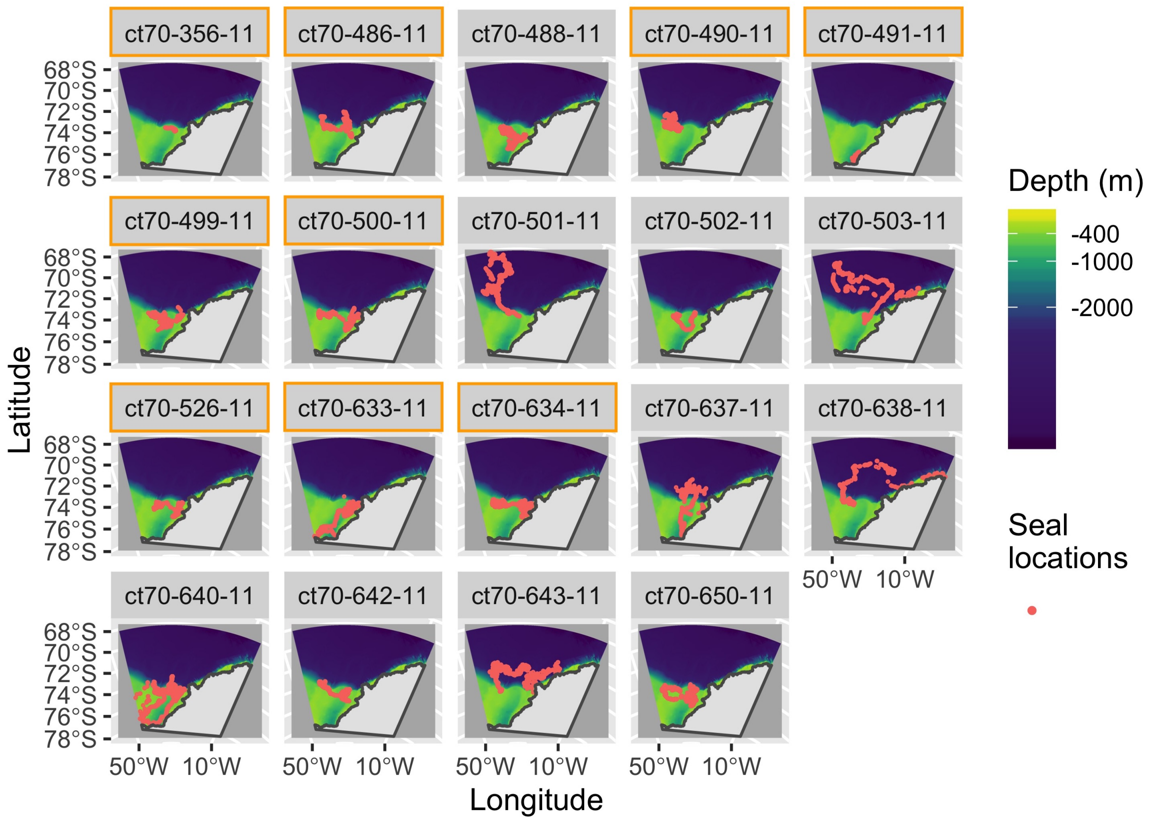


Figure S1. Individual satellite tracks from 19 instrumented Weddell seals carrying CTD-SRDL tags deployed in February 2011. The subplot labels are colour-coded by sex: the plots with a gold border around the heading come from male seals, the rest from females. Dive characteristics are detailed in Table S1. The background colour represents the bathymetry (depth in metres at a 0.5km resolution). The colour bar has been scaled so that yellow areas represent the continental shelf and dark blue areas represent the deep ocean. These are predicted locations from a correlated random walk model fitted to the original data, accounting for the estimated location error provided by CLS Argos, using the `foieGras` package in `R` [2].

### S1.1 Data collection rules

The beginning of a haulout record is triggered when the tag is dry for at least 10min - the animal is considered to be hauled out on ice or land. A surface event is triggered when the tag is wet but there has been no dive for at least 9min - the animal is considered to be in the water but not diving). A dive event is triggered when the tag is wet and the depth is greater than 6m for at least 8sec - the animal is considered to be actively diving. An detailed explanation of how behavioural records are collected by CTD-SRDLs is provided in [3].
