## Supplementary material 2 for "Sex-specific variation in the use of vertical habitat by a resident Antarctic top predator"

### S2.1 Motivation

In contrast to [1] we do not consider the first and last segments (initial descent and final ascent) of each dive. We do this because Weddell seals likely have to travel some distance horizontally under the ice from and to their breathing holes and this may produce shallow-angle swimming unrelated to foraging. The findings of [2] also support the exclusion of "commuting" segments from being important for foraging. This post hoc analysis of the PrCA data from [1] shows that only a small percentage of PrCA behaviours occur in the first and last segments (6%), which is further evidence that we are unlikely to be excluding highly important foraging behaviour by removing them from our analysis.

### S2.2 Background

Heerah *et al.* [3, 4] developed a method for identifying within-dive foraging from time-depth records, such as the dives returned by SMRU CTD-SRDLs abstracted using the broken-stick algorithm ([5, 6]. Dive records returned from CTD-SRDLs consist of six time-depth points - two surface points and four points at depth - identified by a broken-stick algorithm that is implemented on-board the tag [6]. The method presented by Heerah *et al.* [1] is based on the identification of the most sinuous and low speed movement phases of a dive (which they call hunting phases, and are in contrast with straighter and faster movement during vertical transit phases), by transposing the Area Restricted Search concept into the vertical dimension [7, 8]. While the hunting index developed by Heerah *et al.* [1] has been validated using acceleration data for southern elephant seals [3, 4], it is only recently that the authors validated its use for Weddell seals, as 3D acceleration datasets were not previously available (Heerah *et al.* [1], and Table S2.1 below).

| PTT | Mass (kg) | Length (cm) | Deployment date (retrieval) | Transmission duration (days) | Total dives | Dives per day | Maximum dive depth (m) | Dive duration (min) | PrCA (sec) |
|---|---|---|---|---|---|---|---|---|---|
| *143467* | 238 | 227 | 27/11/2014 | 49 | 667 | 13±1 | 120±2 | 11±0.1 | 46±1 |
| *143468* | 436 | 251 | 27/11/2014 (23/01/2015) | 25 | 221 | 9±1 | 120±5 | 14±0.5 | 60±3 |
| *143469* | 339 | 236 | 29/11/2014 | 10 | 55 | 5±1 | 82±6 | 10±1 | 24±4 |
| *143470* | 358 | 255 | 29/11/2014 | 48 | 469 | 10±1 | 129±3 | 15±0.3 | 61±1 |
| *All* | 343±41 | 242±7 | - | 33±9 | 353±135 | 9±2 | 122±1 | 13±0.1 | 55±1 |

Table S2.1 General information on tag deployment and transmission outputs. Data are given for four adult Weddell seals equipped with DSA tags at Dumont D'Urville (66°40' S 140° E) in November 2014. PrCA corresponds to the estimated number of seconds spent in prey capture attempt behaviour by the DSA tag algorithm (See Cox *et al.* [9] and Heerah *et al.* [1] for more details [1]).

To obtain high resolution data, Weddell seals were equipped with a new generation of device, known as a DSA tag (SCOUT-DSA-296 tag, Wildlife Computers; [9]). These were head-mounted on the Weddell seals at Dumont D'Urville, Antarctica. The DSA tag measures 86x85x29mm and weighs 192 g (see also Cox *et al.* [9]). It comprises an Argos transmitter, alongside a pressure sensor (recording rate of 1Hz, resolution of 0.5m and accuracy of ±1m + 1% of a reading), tri-axial accelerometer (recording rate of 16Hz) and a wet-dry sensor. Dives were defined as events that lasted at least 60sec with a maximum depth that exceeded 15m.

The results in Heerah *et al.* [1] show that hunting phases (low vertical speed phases) within a dive, inferred from either archived high-resolution or transmitted low-resolution dive profiles, were associated with most of prey capture attempts (PrCA). This is the major contribution of their work to our study: they found a high correlation between vertical speed and the number of prey capture attempts. This means that even when only low-resolution dive data are available, as is common when tags are deployed in remote areas and cannot be recovered, segments of the dive with low vertical speed can be assumed, with confidence, to be hunting segments.

We investigate the implications of their method for abstracted dives received from SMRU CTD-SRDL tags, used in our study of Weddell seal diving behaviour from the Weddell Sea. We do this by exploring 1) the distribution of transmitted PrCAs within a dive at the dive segment scale, which is the format in which dives are transmitted by SMRU CTD-SRDLs (see main text Methods section for references), and 2) the dive parameters (duration, depth) associated with these hunting segments. Here, we present a post-hoc analysis of the low-resolution dataset analysed in Heerah *et al.* [1], in order to define hunting phases and associated parameters more accurately for abstracted dives returned by CTD-SRDLs.

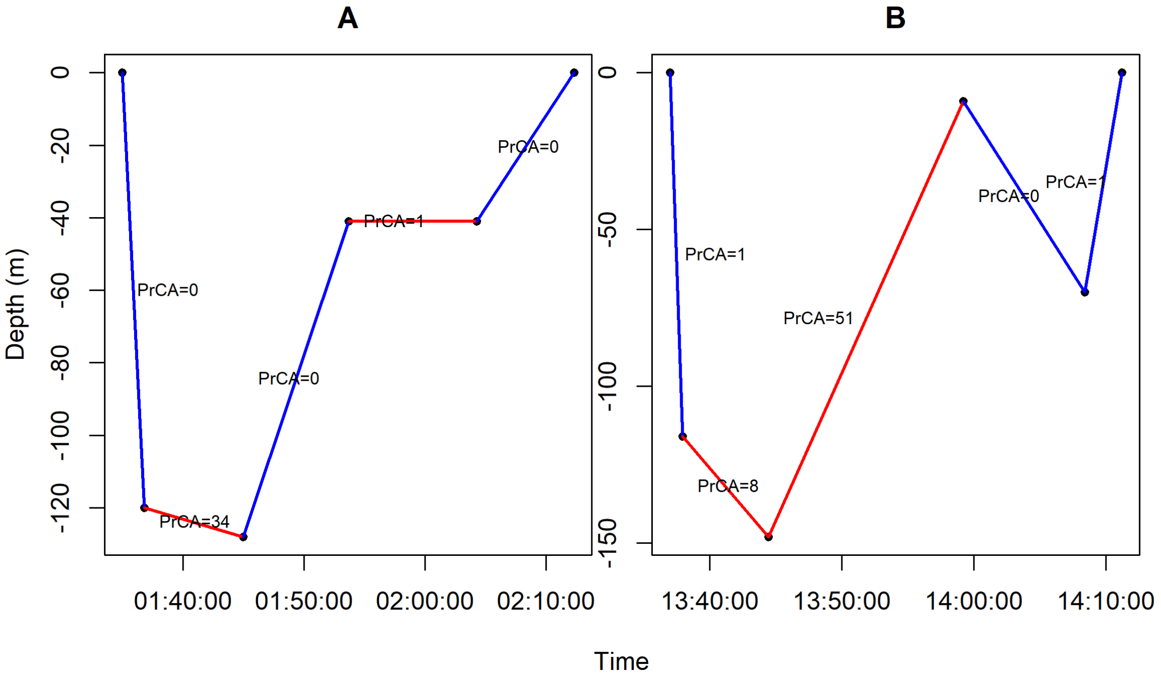


Figure S2.1. Identification of hunting behaviour. Example of two (out of 1412) DSA low-resolution dives that were transmitted, along with the time spent in PrCA behaviours per segment (See Cox *et al.* [9] and Heerah *et al.* [1] for more details – acceleration records were automatically processed on board the DSA tag before transmission). Red lines represent broken stick segments associated with hunting behaviours (segments associated with reduced vertical speed of low-resolution dives, vertical speed $\leq$ 0.5 m/sec). Conversely, blue lines represent segments associated with transit behaviours (segments associated with increased vertical speed of low-resolution dives, vertical speed $>$ 0.5 m/sec) behaviours.

### S2.3 Identification of within-dive foraging effort

#### Hunting index vs. PrCA behaviours estimated from three-axis acceleration

As for CTD-SRDLs, dive data returned by DSA tags comes in an abstracted form, consisting of 6 time points with an associated depth; 2 at the surface at the beginning and end of the dive, and 4 at depth. The 4 points at depth are chosen using a broken-stick algorithm (Cox *et al.* [9], Photopoulou *et al.* [5, 6]). This creates a piecewise-linear time-depth dive profile made up of 5 segments. Each of these segments has an associated start time and start depth, as well as an end time and end depth. In addition, each DSA segment has an associated time spent in PrCA, estimated by an on-board algorithm (Cox *et al.* [9], Heerah *et al.* [1], and Table S2.1).

| Dive segments | Proportion of PrCA occurring within hunting segments | Proportion of hunting segments with PrCA | without PrCA | $R^2$ correlation coefficient: hunting segment duration vs time spent in PrCA behaviour |
|---|---|---|---|---|
| All | 94 % | 83 % | 17 % | 0.76 |
| 1 And 5 | 39 % | 54 % | 46 % | 0.43 |
| 2, 3 and 4 | 98 % | 88 % | 12 % | 0.80 |

Table S2.2. Summary of foraging effort distribution throughout Weddell seal dives, for all individuals pooled together, based on vertical segment speed. We consider the outputs from 1) all segments pooled, 2) the first and last segments (s1 and s5) and 3) the three intermediate segments (s2, s3, and s4) from the transmitted dive profiles. We refer to dive segments with vertical speed lower than 0.5m/sec as hunting segments. We present the percentage of prey capture attempts (PrCA) that take place in hunting segments, the proportion of hunting segments that contain one or more PrCA, the proportion of hunting segments with no PrCA, and the Spearman rank correlation coefficient ($R^2$) between the duration of a hunting segment and the sum of time spent in PrCA during that segment.

Overall, Heerah *et al.* [1] found that low vertical speed (below 0.5m/sec) is associated with a high number of PrCA in Weddell seals (Figure S2.1, shallow angle-low vertical speed segments shown in red, and the number of PrCA annotated on each segment) and use this threshold as an index of hunting behaviour. Our analysis shows that the first and last segments of each dive (s1 and s5) are generally faster (greater vertical speed than expected for hunting behaviour) than the middle three segments (Figure S2.2 A) and therefore more likely to be transiting behaviour. The majority (94%) of observed PrCA behaviours occurred during the three middle segments (Figure S2.2 B, 24 %, 42 % and 28 % for segments 2 to 4, respectively). The remaining 6% of PrCA behaviours occurred in segments 1 and 5 (Figure S2.2 B, 1% and 5% of total PrCAs occurred in segment 1 and 5, respectively).

If we only consider the intermediate dive segments (s2, s3 and s4), 98% of the PrCA behaviours observed within s2 to s4, occurred when vertical speed was low – below the threshold of 0.5 m/sec. This suggests that the 0.5 m/sec threshold is a good proxy for hunting in this middle portion of a dive. This relationship breaks down for the first and last segments. Only 39% of PrCA behaviours observed within s1 and s5, occurred when the vertical speed was less than 0.5m/sec. It follows that the remaining 61% of PrCA occurred when the vertical speed was above the 0.5 m/sec threshold.

These results provide compelling evidence that, when analysing within-dive hunting behaviour, excluding the first and last segments of a dive yields a better correspondence between vertical speed and the number of PrCA (Table S2.2) by reducing the misclassification rate of dive segments as hunting segments. Although it is clear that hunting can occur during the first and last segments, it is not where hunting is concentrated. Low vertical speed during these segments likely comes from the seals exiting or entering the complex under-ice environment associated with breathing holes, since these segments necessarily contain transit behaviour from/to the surface. Based on these results, we only consider the intermediate segments in our analysis, knowing that, if anything, it increases the accuracy of the hunting index in detecting foraging activity, by reducing the false positive rate.

Heerah *et al.* [1] clearly show that hunting can occur at several depths within a dive (also, Figure S2.3 B below). However, for simplicity, we wanted to summarise hunting depth using a single metric for each dive and extract *in situ* environmental variables (temperature and salinity) at that depth to use as covariates on the transition probability matrix in the hidden Markov model. We chose to use the depth of the longest in duration hunting segment. We show that this is a reasonable simplification because there is a high degree of correlation between the depth of the longest in duration hunting segment and the number of PrCA behaviours, in the high-resolution dataset presented in Heerah *et al.* [1] (Figure S2.3 A). In other words, if a segment has a vertical speed lower than 0.5m/sec, the

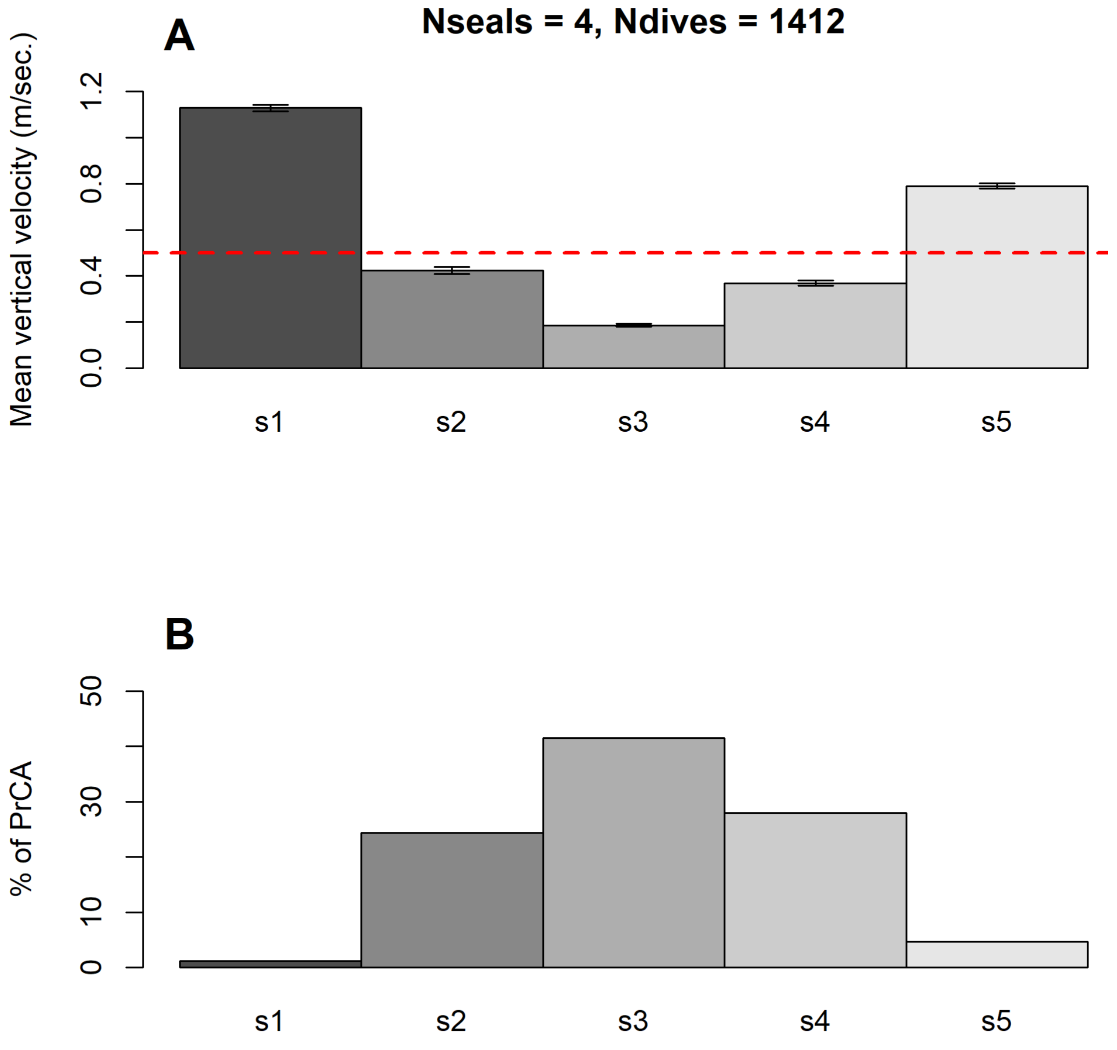


Figure S2.2. Segment-by-segment analysis of dive parameters behaviours displayed as barplots: (A) mean vertical speed (the estimated threshold for hunting behavior is shown as a red dotted line at 0.5m/sec), and (B) percentage of total PrCAs associated with each segment of DSA transmitted dives. On average, segments 1 and 5 (s1 and s5) are associated with transit behaviour (A, vertical speed $>$ 0.5 m/sec). It follows that these segments are associated with a small percentage of PrCA occurrences (B, 1% and 5 % of total PrCAs occurred in segment 1 and 5, respectively). In contrast, segments 2 to 4 are, on average, associated with hunting behaviour (A, vertical speed $\leq$ 0.5 m/sec). 94% of total PrCAs occurred in these segments (B, 24 %, 42 % and 28 % for segments 2 to 4 respectively)

longer the duration of the segment the greater the number of PrCA it contains. Again, this relationship held between the duration of hunting segments and the time spent in PrCA behaviours for all segments together and for the three middle segments, but less so for the first and last segments (Table S2.1).

A comparison of the maximum dive depth and the depth of the longest hunting segment shows that there is a difference between the two (Figure S2.3 B) and it makes it worth considering environmental conditions at the depth of the longest hunting segment, as we have done in our analyses.

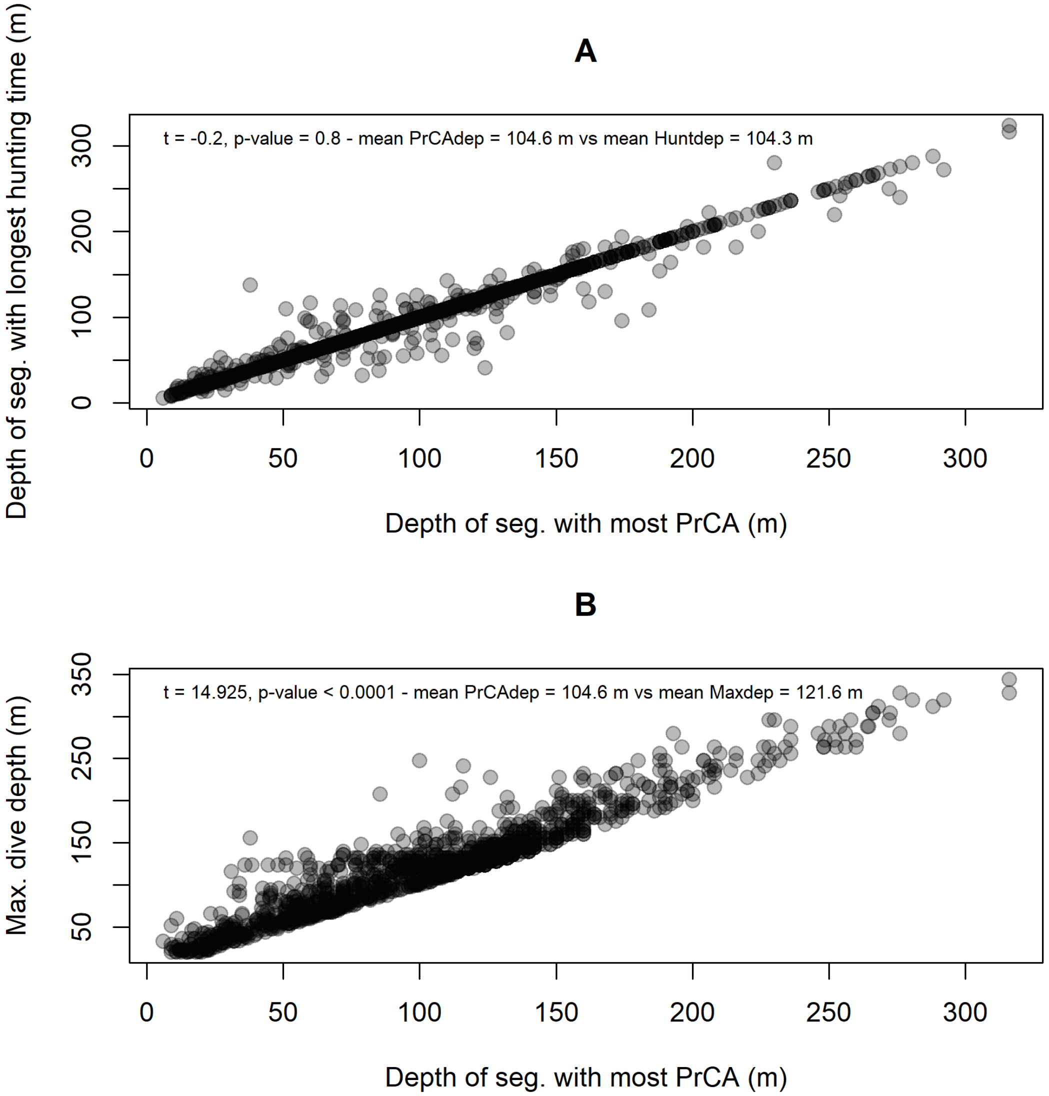


Figure S2.3. Occurrences of foraging at depth. We compare segment depth where most PrCA occurred, with the depth of the longest hunting segment (A, vertical speed $\leq$ 0.5 m/sec) and maximum dive depth (B). In both cases, compared depths were highly correlated (Spearman correlation, $R^2$=0.92). However, statistical comparison of the mean depths, indicated significant differences between the depths of the segments where most PrCA behaviours occurred and maximum dive depth (B). On average, the maximum dive depth is deeper than the depths where most PrCA behaviours occurred (B). No significant differences were observed between the depths of the longest hunting segment and the depths where most PrCA behaviours occurred (A).
