## Supplementary material 3 for "Sex-specific variation in the use of vertical habitat by a resident Antarctic top predator"

### S3.1 Matching CTD data to behavioural data

Not every dive profile (time-depth trace) had an associated CTD profile (temperature-salinity-depth trace), so we interpolated practical salinity (hereafter, salinity) linearly in time, between time points where CTD data were available, and assigned it to the intervening dive times. We are most interested in the oceanographic conditions encountered by seals at depths where they are foraging. To link hunting behaviour with *in situ* oceanographic conditions we calculate the mean depth of the longest in duration hunting segment in each dive (dives from CTD-SRDLs are made up of 5 segments, [1, 2]) and extract salinity at that depth. For the small number of dives with no detectable hunting segments there is no depth at which to extract salinity and these values are missing. For surface events we extracted surface variables whereas for haulout events, where the seal is not in the water, we use the conditions experienced during the most recent previous dive.

To obtain a value for salinity at the relevant depth in each dive (hunting depth) and surface event (shallowest depth), we interpolated the hydrographic data from each seal separately at 10m intervals between the surface (5m) and the deepest hunting depth in the whole dataset (805m). For each row in the behavioural data, we were then able to chose the interpolated salinity at the depth that was closest to the depth of interest.

We carried out the interpolation using a weighted interpolation function (`wtd.quantile()` from the R package `Hmisc`). We derived the weights by iteratively evaluating the density of a normal distribution at the CTD profile timestamps. At each iteration the normal distribution was centred on a dive timestamp, with a standard deviation equal to the bandwidth. This results in a vector of densities corresponding to how close the CTD timestamps were to the dive timestamp, with nearby (in time) CTD profiles having greater densities. We chose a 6hr bandwidth so that it would include at least one CTD profile, since the tags were programmed to collect at least one CTD profile every 4hrs. The resulting weights convey that CTD profiles closer in time to a dive, for which we wish to obtain interpolated values, are more relevant. We use these weights to assign median values (value at the 0.5 quantile) of salinity to each of the dives (weighted interpolation). We only used CTD profiles scoring 1 on the Argo quality control flag scale, where 1 flags "Good data" [3].

### S3.2 Extracting bathymetry data at behaviour locations

There is no explicitly spatial component to our analysis of seal behaviour, however, an estimate of bathymetry - a spatial field - was required for each behaviour location to extract one of the state-dependent variables: proportion of bathymetry reached. Locations obtained via Argos CLS [4] are prone to large errors so we fitted a state-space model to the seal locations using the R package `foieGras` [5] prior to matching them with bathymetry. This approach takes into account the degree of error associated with each location, as estimated by Argos CLS, and predicts the most likely trajectory. We used the IBCSO bathymetric dataset [6] and extracted values at the model-filtered locations of seal behaviours using the `extract` function from the `raster` R package. The IBCSO bathymetry dataset has a spatial resolution of 500x500m.

### S3.3 A note on the state-dependent variable: proportion of bathymetry reached

Hunting depth and proportion of bathymetry reached are both biologically informative variables but are highly correlated in this dataset. Including correlated variables as state variables in an HMM violates the assumption of longitudinal conditional independence that we use in our analysis to simplify the modelling: the random vectors that make up the state variables are assumed to be mutually independent given the states [7]. To get around this we include the proportion of bathymetry reached as an indicator variable for identifying benthic dives. Instead of estimating distributional parameters for each state, we estimate the probability that dives of a given state are benthic, according to a threshold value (0.80 of bathymetry) obtained from the literature ([8, 9] and references therein).

[3] Argo Data Management Team. Argo user's manual V3.3; 2019.

[4] Argos. Argos User's Manual:Worldwide tracking and environmental monitoring by satellite; 2016.

[5] Jonsen I, Patterson T. foieGras: Fit Continuous-Time State-Space and Latent Variable Models for Filtering Argos Satellite (and Other) Telemetry Data and Estimating Movement Behaviour. CRAN; 2019. R package version 0.4.01. Available from: `https://cran.r-project.org/package=foieGras`.

[6] Arndt JE, Schenke HW, Jakobsson M, Nitsche FO, Buys G, Goleby B, et al. The International Bathymetric Chart of the Southern Ocean (IBCSO) Version 1.0—A new bathymetric compilation covering circum-Antarctic waters. Geophysical Research Letters. 2013;40(12):3111–3117.

[7] Zucchini W, MacDonald IL, Langrock R. Hidden Markov models for time series: an introduction using R. Chapman and Hall/CRC; 2017.

[8] Tremblay Y, Cherel Y. Benthic and pelagic dives: a new foraging behaviour in rockhopper penguins. Marine Ecology Progress Series. 2000;204:257–267.

[9] Heerah K, Andrews-Goff V, Williams G, Sultan E, Hindell M, Patterson T, et al. Ecology of Weddell seals during winter: Influence of environmental parameters on their foraging behaviour. Deep Sea Research Part II: Topical Studies in Oceanography. 2013;88-89:23 – 33.
