## Supplementary material 4 for "Sex-specific variation in the use of vertical habitat by a resident Antarctic top predator"

### S4.1 Model structure and implementation

A partially unobserved Markov chain is assumed to determine the behavioural states and the parameters of the state-dependent distributions associated with the observed variables. While the type of event (haulout, surface and dive) is known from the data, dives are not further distinguishable into types or classes from the data alone. In this HMM analysis, surface and haulout states are threrefore treated as known, while the dive states are treated as unknown states and our research question relates to inferring information about the different types of dives carried out by Weddell seals. We fit the model to all seals of each sex jointly and therefore estimate one set of parameters for females and one for males. We explored fitting a single model to all seals but this model structure did not allow us to detect differences in diving behaviour and compare the effects of the covariates in females and males. By the time one has made all parameters sex-dependent it becomes more practical to fit separate models.

The HMM parameters are estimated using numerical maximisation of the likelihood, implemented in R [1], using the `nlm` function, with the computation of the covariate-dependent transition probability matrices and the forward algorithm coded in `C++`. The forward algorithm is an efficient way of evaluating the likelihood and is one reason for the popularity of HMMs – it makes them relatively fast to fit. It corresponds to a recursive calculation of the likelihood with computational costs only linear in the number of observed time points and renders numerical maximum likelihood estimation feasible [2].

Based on prior knowledge about Weddell seals' diving behaviour from the literature, and the dive records from the current dataset, we fitted HMMs with 3 or 4 diving states (i.e. 5- and 6-state HMMs, considering both dive and non-dive states). For the male dataset, the 5-state HMM fitted the data reasonable well, while the 6-state model suffered from numerical instabilities and was difficult to interpret. For the female dataset, however, the 6-state model was stable and better able to capture the more variable dive types carried out in shallow and deep water by female seals. Using a different number of dive states for females and males corresponds well with the fact that the female seals in our dataset tend to use deep water regions more, while male seals seem to avoid them (as illustrated in Figure 1, main manuscript). Very deep water does not allow for benthic dives, so we assume that diving behaviour might be different to coastal and continental shelf areas. Based on the above considerations, we present the results of the 5-state HMM for males, and 6-state HMM for females [3].

### S4.2 Likelihood of the HMMs

The likelihood of an HMM with N states and observation vectors $z_1, ..., z_T$ can be written as a matrix product:

$$L = \delta \mathbf{P}(\mathbf{z_1}) \prod_{t=2}^{T} \Gamma_t \mathbf{P}(\mathbf{z_t}) 1' \qquad (1)$$

where $\delta$ is a row-vector containing the initial state distribution, $\Gamma_t$ represents the $N \times N$ transition probability matrix at time point t and 1 is a row-vector of ones. $P$ denotes a $N \times N$ diagonal matrix containing the values of the $N$ joint state-dependent densities evaluated at the observation vector

$\mathbf{z_t}$. We assume the observed variables to be contemporaneously conditionally independent, given the current state. Thus, for each state, the joint state-dependent density is the product of the univariate state-dependent densities which are associated to the observed variables.

In this analysis, $\mathbf{z_t}$ corresponds to the vector: duration, hunting depth, proportion of dive time spent hunting, proportion of bathymetry, and salinity at hunting depth, observed at time $t$ given that a dive event occurred. For haulout and surface events, $\mathbf{z_t}$ only contains the observed duration.

To investigate the influence of time of day and season on the diving behaviour, temporal covariates are incorporated into the transition probabilities using multinomial logit links (with the probability to remain in the state as the reference category):

$$\ln\left(\frac{\gamma_{ij}(t)}{\gamma_{ii}(t)}\right) = \beta_{0ij} + \beta_{1ij}\cos(2\pi \mathrm{hr}_t/24) + \beta_{2ij}\sin(2\pi \mathrm{hr}_t/24) + \beta_{3ij}\mathrm{week}_t + \beta_{4ij}\cos(2\pi \mathrm{hr}_t/24)\mathrm{week}_t + \beta_{5ij}\sin(2\pi \mathrm{hr}_t/24)\mathrm{week}_t \quad (2)$$

where $\gamma_{ij}(t)$ denotes the probability of switching from state $i$ to state $j$ at time $t$. Given we are not fitting any random effects, the log-likelihood of interest is the sum of log-likelihoods corresponding to the different seals within each sex.
