## Supplementary material 5 for "Sex-specific variation in the use of vertical habitat by a resident Antarctic top predator"

#### S5.1 Transition probability matrices

We present transition probability matrices for the conditions presented in Figure 3 of the main text. We provide point estimates and 95% confidence intervals for the probability of transitioning between states, at two times of day (midnight and midday), during three weeks of the year (week 7, week 15 and week 24), for female and male Weddell seals. The estimates on the diagonal of each transition probability matrix are shown in **bold**.

These results suggest persistence in behaviour - once a seal enters a movement state it tends to remain it in - but the degree of persistence varies substantially between night and day, seasonally and between the two sexes. In terms of behavioural succession, there is a clear overall trend of regular switching between haulout and surface behaviour, and a tendency to switch between adjacent states in order of depth.

It is not fair to directly compare transition probabilities for diving states between males and females due to the different number of states, but meaningful comparisons can still be made.

In general, in all weeks and in both sexes, benthic dives have a higher persistence in the day. This is also true for pelagic dives (state present in females only) at least up to week 15. By week 24, persistence in state is similar for female pelagic and benthic dives.

##### S5.1.1 Females

In week 7 (14-20 February 2011) there is little difference in light level between midday and midnight, but female Weddell seals show diurnal variation in state transition probabilities all the same. At midnight, females are most likely to switch to shallower diving states, and they are most likely to continue to engage in epipelagic and pelagic diving (Table S5.1). At midday, diving behaviour moves away from shallower diving and into the two deeper dive states, pelagic and benthic diving. Surface events seem to be a stepping stone for entering benthic diving, while switches into pelagic diving are more likely from epipelagic diving. Non-diving behaviours have a lower degree of residency at midday.

Females - WEEK 7

| MIDNIGHT | *To state* | | | | | |
|---|---|---|---|---|---|---|
| *From state* | Haulout | Surface | Shallow dive | Epipelagic dive | Pelagic dive | Benthic dive |
| Haulout | **0.40 (0.32-0.47)** | 0.45 (0.34-0.55) | 0.08 (0.05-0.14) | 0.06 (0.03-0.10) | 0.01 (0.01-0.02) | 0.00 (0.00-0.01) |
| Surface | 0.24 (0.14-0.37) | **0.34 (0.27-0.41)** | 0.19 (0.11-0.32) | 0.11 (0.05-0.22) | 0.07 (0.03-0.14) | 0.05 (0.02-0.13) |
| Shallow dive | 0.02 (0.01-0.03) | 0.13 (0.07-0.21) | **0.42 (0.34-0.50)** | 0.22 (0.13-0.34) | 0.20 (0.11-0.33) | 0.02 (0.01-0.06) |
| Epipelagic dive | 0.02 (0.01-0.02) | 0.06 (0.04-0.11) | 0.16 (0.09-0.27) | **0.54 (0.47-0.62)** | 0.22 (0.13-0.34) | 0.00 (0.00-0.00) |
| Pelagic dive | 0.02 (0.01-0.03) | 0.12 (0.07-0.20) | 0.09 (0.05-0.16) | 0.13 (0.07-0.24) | **0.63 (0.55-0.70)** | 0.00 (0.00-0.03) |
| Benthic dive | 0.02 (0.01-0.04) | 0.23 (0.12-0.38) | 0.16 (0.07-0.31) | 0.14 (0.07-0.28) | 0.13 (0.05-0.28) | **0.32 (0.25-0.40)** |
| MIDDAY | Haulout | Surface | Shallow dive | Epipelagic dive | Pelagic dive | Benthic dive |
| Haulout | **0.21 (0.16-0.25)** | 0.41 (0.30-0.54) | 0.06 (0.04-0.10) | 0.03 (0.02-0.05) | 0.18 (0.11-0.27) | 0.12 (0.06-0.23) |
| Surface | 0.19 (0.10-0.31) | **0.32 (0.24-0.41)** | 0.09 (0.05-0.17) | 0.04 (0.02-0.09) | 0.13 (0.07-0.24) | 0.23 (0.10-0.45) |
| Shallow dive | 0.07 (0.04-0.12) | 0.13 (0.08-0.21) | **0.55 (0.46-0.63)** | 0.07 (0.04-0.11) | 0.07 (0.04-0.12) | 0.12 (0.05-0.26) |
| Epipelagic dive | 0.02 (0.01-0.03) | 0.05 (0.03-0.08) | 0.13 (0.07-0.23) | **0.37 (0.30-0.45)** | 0.30 (0.19-0.43) | 0.13 (0.07-0.25) |
| Pelagic dive | 0.02 (0.01-0.03) | 0.06 (0.04-0.11) | 0.03 (0.02-0.05) | 0.04 (0.02-0.08) | **0.80 (0.74-0.85)** | 0.04 (0.02-0.11) |
| Benthic dive | 0.02 (0.01-0.04) | 0.06 (0.03-0.11) | 0.06 (0.03-0.12) | 0.02 (0.01-0.04) | 0.03 (0.01-0.08) | **0.81 (0.75-0.86)** |

Table S5.1. State transition probabilities (with 95% confidence intervals) for female Weddell seals in week 7 of 2011 (14-20 February) at midnight (top matrix) and midday (bottom matrix).

In week 15 (11-17 April 2011) nighttime transition probabilities do not seem very different from week 7, except for an increase in probability of continuing to dive epipelagically and a slight decrease in continuing to dive pelagically. There is also a shift in daytime diving with a decrease in the probability of continued pelagic diving behaviour, compared to week 7 (Table S5.2).

Females - WEEK 15

| MIDNIGHT | | | *To state* | | | |
|---|---|---|---|---|---|---|
| *From state* | Haulout | Surface | Shallow dive | Epipelagic dive | Pelagic dive | Benthic dive |
| Haulout | **0.39 (0.35-0.44)** | 0.43 (0.36-0.50) | 0.08 (0.07-0.10) | 0.07 (0.05-0.11) | 0.02 (0.01-0.02) | 0.00 (0.00-0.01) |
| Surface | 0.29 (0.22-0.37) | **0.40 (0.35-0.46)** | 0.14 (0.11-0.19) | 0.14 (0.09-0.22) | 0.02 (0.01-0.04) | 0.01 (0.00-0.01) |
| Shallow dive | 0.03 (0.02-0.04) | 0.14 (0.11-0.18) | **0.47 (0.42-0.53)** | 0.32 (0.25-0.40) | 0.02 (0.01-0.04) | 0.01 (0.01-0.02) |
| Epipelagic dive | 0.02 (0.02-0.03) | 0.08 (0.07-0.10) | 0.19 (0.12-0.28) | **0.69 (0.62-0.76)** | 0.01 (0.01-0.02) | 0.00 (0.00-0.01) |
| Pelagic dive | 0.04 (0.03-0.05) | 0.14 (0.11-0.18) | 0.09 (0.06-0.13) | 0.15 (0.10-0.24) | **0.57 (0.51-0.62)** | 0.01 (0.01-0.02) |
| Benthic dive | 0.04 (0.03-0.05) | 0.15 (0.12-0.20) | 0.15 (0.09-0.26) | 0.20 (0.12-0.31) | 0.12 (0.07-0.21) | **0.33 (0.28-0.39)** |
| MIDDAY | | | | | | |
| | Haulout | Surface | Shallow dive | Epipelagic dive | Pelagic dive | Benthic dive |
| Haulout | **0.27 (0.24-0.30)** | 0.44 (0.38-0.50) | 0.06 (0.05-0.07) | 0.04 (0.03-0.06) | 0.10 (0.07-0.15) | 0.09 (0.05-0.15) |
| Surface | 0.24 (0.19-0.30) | **0.39 (0.35-0.44)** | 0.09 (0.07-0.11) | 0.05 (0.03-0.09) | 0.11 (0.07-0.17) | 0.12 (0.06-0.21) |
| Shallow dive | 0.07 (0.05-0.08) | 0.15 (0.13-0.19) | **0.45 (0.41-0.49)** | 0.14 (0.10-0.18) | 0.11 (0.07-0.16) | 0.08 (0.05-0.15) |
| Epipelagic dive | 0.04 (0.03-0.05) | 0.09 (0.07-0.10) | 0.12 (0.07-0.20) | **0.42 (0.37-0.47)** | 0.23 (0.16-0.33) | 0.10 (0.06-0.16) |
| Pelagic dive | 0.04 (0.03-0.05) | 0.06 (0.05-0.07) | 0.04 (0.03-0.06) | 0.07 (0.05-0.12) | **0.71 (0.65-0.75)** | 0.08 (0.05-0.15) |
| Benthic dive | 0.02 (0.01-0.02) | 0.04 (0.03-0.05) | 0.03 (0.02-0.06) | 0.05 (0.03-0.08) | 0.07 (0.04-0.12) | **0.79 (0.74-0.83)** |

Table S5.2. State transition probabilities (with 95% confidence intervals) for female Weddell seals in week 15 of 2011 (11-17 April) at midnight (top matrix) and midday (bottom matrix).

In week 24 (13-19 June 2011), the persistence of nighttime epipelagic dives increses further, compared with week 15. In the day, deep diving becomes more variable, with lower probabilities of continuing to dive pelagically and benthically. Nonetheless, these two dive states still have the highest residency probabilities. At midday switching behaviour overall seems to increase (off-diagonal elements) (Table S5.3).

Females - WEEK 24

| MIDNIGHT | | | *To state* | | | |
|---|---|---|---|---|---|---|
| *From state* | Haulout | Surface | Shallow dive | Epipelagic dive | Pelagic dive | Benthic dive |
| Haulout | **0.39 (0.30-0.49)** | 0.41 (0.28-0.55) | 0.08 (0.05-0.14) | 0.09 (0.05-0.17) | 0.02 (0.01-0.04) | 0.00 (0.00-0.01) |
| Surface | 0.32 (0.17-0.51) | **0.43 (0.31-0.56)** | 0.09 (0.04-0.18) | 0.16 (0.07-0.31) | 0.01 (0.00-0.02) | 0.00 (0.00-0.00) |
| Shallow dive | 0.05 (0.02-0.10) | 0.12 (0.06-0.21) | **0.43 (0.32-0.55)** | 0.40 (0.26-0.56) | 0.00 (0.00-0.00) | 0.00 (0.00-0.01) |
| Epipelagic dive | 0.03 (0.02-0.05) | 0.08 (0.04-0.14) | 0.18 (0.10-0.30) | **0.70 (0.59-0.78)** | 0.00 (0.00-0.00) | 0.02 (0.01-0.04) |
| Pelagic dive | 0.08 (0.04-0.14) | 0.17 (0.09-0.30) | 0.08 (0.04-0.16) | 0.18 (0.09-0.32) | **0.48 (0.39-0.58)** | 0.01 (0.00-0.03) |
| Benthic dive | 0.07 (0.04-0.13) | 0.09 (0.04-0.20) | 0.14 (0.06-0.28) | 0.27 (0.14-0.46) | 0.10 (0.04-0.26) | **0.33 (0.24-0.43)** |
| MIDDAY | | | | | | |
| | Haulout | Surface | Shallow dive | Epipelagic dive | Pelagic dive | Benthic dive |
| Haulout | **0.34 (0.27-0.42)** | 0.44 (0.32-0.56) | 0.05 (0.03-0.09) | 0.05 (0.03-0.09) | 0.05 (0.03-0.09) | 0.06 (0.03-0.13) |
| Surface | 0.29 (0.17-0.45) | **0.45 (0.36-0.55)** | 0.07 (0.04-0.15) | 0.06 (0.03-0.12) | 0.08 (0.04-0.15) | 0.05 (0.02-0.14) |
| Shallow dive | 0.05 (0.03-0.09) | 0.17 (0.09-0.28) | **0.31 (0.24-0.38)** | 0.26 (0.16-0.39) | 0.17 (0.09-0.28) | 0.05 (0.02-0.13) |
| Epipelagic dive | 0.09 (0.06-0.14) | 0.16 (0.09-0.26) | 0.10 (0.05-0.18) | **0.43 (0.37-0.50)** | 0.16 (0.09-0.26) | 0.06 (0.03-0.12) |
| Pelagic dive | 0.06 (0.04-0.11) | 0.05 (0.03-0.10) | 0.05 (0.03-0.09) | 0.12 (0.06-0.22) | **0.56 (0.45-0.66)** | 0.15 (0.06-0.35) |
| Benthic dive | 0.1 (0.01-0.02) | 0.03 (0.01-0.06) | 0.01 (0.01-0.03) | 0.12 (0.06-0.22) | 0.15 (0.06-0.33) | **0.68 (0.55-0.79)** |

Table S5.3. State transition probabilities (with 95% confidence intervals) for female Weddell seals in week 24 of 2011 (13-19 June) at midnight (top matrix) and midday (bottom matrix).

#### S5.1.2 Males

In week 7 (14-20 February 2011) male Weddell seals are more likely than females to remain in the shallow diving state and the benthic diving state, at the same time of year. During the day, males have a high probability of continued shallow diving and benthic diving, and residency epipelagic diving becomes less likely (Table S5.4).

Males - WEEK 7

| MIDNIGHT | | | *To state* | | | |
|---|---|---|---|---|---|---|
| *From state* | Haulout | Surface | Shallow dive | Epipelagic dive | Pelagic dive | Benthic dive |
| Haulout | **0.28 (0.22-0.35)** | 0.46 (0.34-0.58) | 0.19 (0.12-0.28) | 0.05 (0.03-0.08) | | 0.03 (0.01-0.06) |
| Surface | 0.17 (0.11-0.26) | **0.46 (0.38-0.54)** | 0.26 (0.16-0.40) | 0.06 (0.03-0.10) | | 0.05 (0.05-0.09) |
| Shallow dive | 0.03 (0.02-0.05) | 0.13 (0.08-0.21) | **0.65 (0.58-0.72)** | 0.14 (0.09-0.20) | | 0.05 (0.03-0.08) |
| Epipelagic dive | 0.01 (0.01-0.02) | 0.11 (0.06-0.19) | 0.22 (0.14-0.32) | **0.63 (0.55-0.71)** | | 0.03 (0.01-0.05) |
| Pelagic dive | | | | | | |
| Benthic dive | 0.02 (0.01-0.04) | 0.14 (0.07-0.25) | 0.11 (0.06-0.21) | 0.21 (0.12-0.35) | | **0.52 (0.42-0.61)** |
| MIDDAY | | | | | | |
| | Haulout | Surface | Shallow dive | Epipelagic dive | Pelagic dive | Benthic dive |
| Haulout | **0.21 (0.17-0.26)** | 0.37 (0.25-0.51) | 0.15 (0.10-0.22) | 0.15 (0.10-0.23) | | 0.11 (0.06-0.21) |
| Surface | 0.27 (0.18-0.39) | **0.33 (0.26-0.41)** | 0.10 (0.06-0.17) | 0.04 (0.02-0.08) | | 0.26 (0.16-0.40) |
| Shallow dive | 0.08 (0.05-0.12) | 0.09 (0.06-0.14) | **0.66 (0.60-0.71)** | 0.07 (0.04-0.10) | | 0.10 (0.07-0.16) |
| Epipelagic dive | 0.09 (0.06-0.14) | 0.04 (0.02-0.07) | 0.20 (0.13-0.30) | **0.44 (0.37-0.52)** | | 0.23 (0.14-0.34) |
| Pelagic dive | | | | | | |
| Benthic dive | 0.02 (0.01-0.03) | 0.05 (0.02-0.09) | 0.07 (0.04-0.13) | 0.03 (0.01-0.04) | | **0.84 (0.79-0.89)** |

Table S5.4. State transition probabilities (with 95% confidence intervals) for male Weddell seals in week 7 of 2011 (14-20 February) at midnight (top matrix) and midday (bottom matrix).

In week 15 (11-17 April 2011), nighttime transition probabilities show higher residency in shallow diving behaviour than females at the same time of year. The clearest differences between night and daytime transition probabilities for males at this time of year are a reduction in the probability of continuing to dive epipelagically, and the consistently high probability of remaining in a benthic diving state in the day (Table S5.5).

Males - WEEK 15

| MIDNIGHT | | | *To state* | | | |
|---|---|---|---|---|---|---|
| *From state* | Haulout | Surface | Shallow dive | Epipelagic dive | Pelagic dive | Benthic dive |
| Haulout | **0.37 (0.32-0.42)** | 0.42 (0.35-0.50) | 0.14 (0.11-0.18) | 0.05 (0.04-0.07) | | 0.01 (0.01-0.02) |
| Surface | 0.22 (0.18-0.27) | **0.49 (0.45-0.54)** | 0.20 (0.15-0.25) | 0.08 (0.05-0.12) | | 0.01 (0.01-0.02) |
| Shallow dive | 0.05 (0.03-0.06) | 0.13 (0.10-0.16) | **0.65 (0.62-0.68)** | 0.16 (0.13-0.19) | | 0.02 (0.01-0.03) |
| Epipelagic dive | 0.03 (0.02-0.04) | 0.09 (0.07-0.11) | 0.24 (0.20-0.28) | **0.61 (0.57-0.65)** | | 0.04 (0.03-0.05) |
| Pelagic dive | | | | | | |
| Benthic dive | 0.06 (0.04-0.10) | 0.14 (0.09-0.21) | 0.19 (0.14-0.24) | 0.29 (0.22-0.37) | | **0.32 (0.28-0.37)** |
| MIDDAY | | | | | | |
| | Haulout | Surface | Shallow dive | Epipelagic dive | Pelagic dive | Benthic dive |
| Haulout | **0.30 (0.27-0.32)** | 0.36 (0.31-0.41) | 0.13 (0.10-0.16) | 0.11 (0.08-0.13) | | 0.11 (0.09-0.14) |
| Surface | 0.23 (0.19-0.27) | **0.40 (0.36-0.45)** | 0.12 (0.09-0.15) | 0.05 (0.03-0.07) | | 0.21 (0.14-0.30) |
| Shallow dive | 0.04 (0.03-0.05) | 0.09 (0.07-0.11) | **0.65 (0.63-0.68)** | 0.12 (0.11-0.14) | | 0.10 (0.07-0.13) |
| Epipelagic dive | 0.07 (0.05-0.09) | 0.07 (0.05-0.08) | 0.18 (0.15-0.21) | **0.51 (0.47-0.54)** | | 0.18 (0.14-0.24) |
| Pelagic dive | | | | | | |
| Benthic dive | 0.02 (0.01-0.03) | 0.05 (0.03-0.07) | 0.06 (0.05-0.08) | 0.06 (0.04-0.08) | | **0.82 (0.79-0.84)** |

Table S5.5. State transition probabilities (with 95% confidence intervals) for male Weddell seals in week 15 of 2011 (11-17 April) at midnight (top matrix) and midday (bottom matrix).

In week 24 (13-19 June 2011), male seals more frequently switch into epipelagic diving behaviour at night compared to week 15, especially from benthic diving. However, benthic diving itself has the lowest residency probability for males at night in week 24. In the day this is reversed, and benthic diving has a high residence probability. This is now almost the only difference between night and day for male seals (Table S5.6). Transitions from all behaviours into benthic diving are higher in the day. There is a clear increase in the probability of remaining in the non-diving states as the season progresses, and this effect is more pronounced in males than in females.

Males - WEEK 24

| MIDNIGHT | | | | | | |
|---|---|---|---|---|---|---|
| | *To state* | | | | | |
| *From state* | Haulout | Surface | Shallow dive | Epipelagic dive | Pelagic dive | Benthic dive |
| Haulout | **0.48 (0.36-0.59)** | 0.37 (0.23-0.53) | 0.10 (0.06-0.15) | 0.06 (0.03-0.09) | | 0.01 (0.00-0.02) |
| Surface | 0.27 (0.16-0.41) | **0.49 (0.39-0.59)** | 0.13 (0.07-0.23) | 0.10 (0.05-0.21) | | 0.00 (0.00-0.01) |
| Shallow dive | 0.07 (0.04-0.12) | 0.12 (0.07-0.21) | **0.63 (0.54-0.71)** | 0.17 (0.11-0.27) | | 0.01 (0.00-0.01) |
| Epipelagic dive | 0.06 (0.03-0.11) | 0.07 (0.03-0.13) | 0.25 (0.15-0.39) | **0.57 (0.47-0.66)** | | 0.06 (0.03-0.11) |
| Pelagic dive | | | | | | |
| Benthic dive | 0.16 (0.08-0.30) | 0.12 (0.05-0.24) | 0.25 (0.13-0.43) | 0.32 (0.17-0.52) | | **0.15 (0.11-0.21)** |
| MIDDAY | | | | | | |
| | Haulout | Surface | Shallow dive | Epipelagic dive | Pelagic dive | Benthic dive |
| Haulout | **0.41 (0.33-0.50)** | 0.33 (0.21-0.46) | 0.10 (0.06-0.15) | 0.06 (0.04-0.10) | | 0.10 (0.05-0.20) |
| Surface | 0.18 (0.11-0.28) | **0.48 (0.40-0.56)** | 0.13 (0.08-0.21) | 0.05 (0.03-0.10) | | 0.16 (0.09-0.27) |
| Shallow dive | 0.1 (0.01-0.02) | 0.08 (0.05-0.13) | **0.60 (0.52-0.68)** | 0.22 (0.15-0.33) | | 0.08 (0.05-0.13) |
| Epipelagic dive | 0.04 (0.03-0.07) | 0.11 (0.06-0.19) | 0.15 (0.09-0.23) | **0.56 (0.48-0.64)** | | 0.14 (0.08-0.22) |
| Pelagic dive | | | | | | |
| Benthic dive | 0.02 (0.01-0.03) | 0.04 (0.02-0.09) | 0.05 (0.02-0.09) | 0.14 (0.08-0.23) | | **0.75 (0.67-0.82)** |

Table S5.6. State transition probabilities (with 95% confidence intervals) for male Weddell seals in week 24 of 2011 (13-19 June) at midnight (top matrix) and midday (bottom matrix).

### S5.2 Model checking

To check if the model fit adequately we 1) computed the pseudo-residuals (conditioned on being in one of the diving states), and 2) simulated data from the model and compared the distribution of simulated state variables to the observed distributions.

#### S5.2.1 Pseudo-residuals

We computed forecast pseudo-residuals (conditional on being in one of the diving states) for each of the observed variables using the approach described in Zucchini *et al.* [1]. We show the histograms of the pseudo-residuals for males and females in Figure S5.1, none of which suggest a systematic lack of fit in the model. Figure S5.2 displays the autocorrelation functions of the pseudo-residuals. Some autocorrelation remained in the residuals for all state-dependent variables. However, the remaining serial correlation is only high for salinity, which might partly be caused by the additional spatial correlation of this variable. For the state classification, duration and depth are the most important, as their corresponding state-dependent distributions are well separated and they describe the key features of the different dive types. For these variables, the remaining autocorrelation is quite low. Thus, although the HMM is not able to capture the full autocorrelation structure in the data, the overall goodness of fit of the pseudo-residuals is satisfactory.

#### S5.2.2 Simulating from the model

Simulating data from the fitted model and comparing it to the observed data is another powerful tool for assessing the models' fit to the data. We show histograms of the observed and simulated data (Figure S5.3 and S5.4, respectively) for dive states with the component distributions colour-coded by state, following the colours in the main text. The state labels for non-diving states (haulout and surface events) are given as known in the model so these distributions are of less interest. The marginal distributions are shown in orange.

There are some small discrepancies in the observed data distributions and the simulated distributions but overall the model structure we have used seems to adequately capture the patterns in the data.

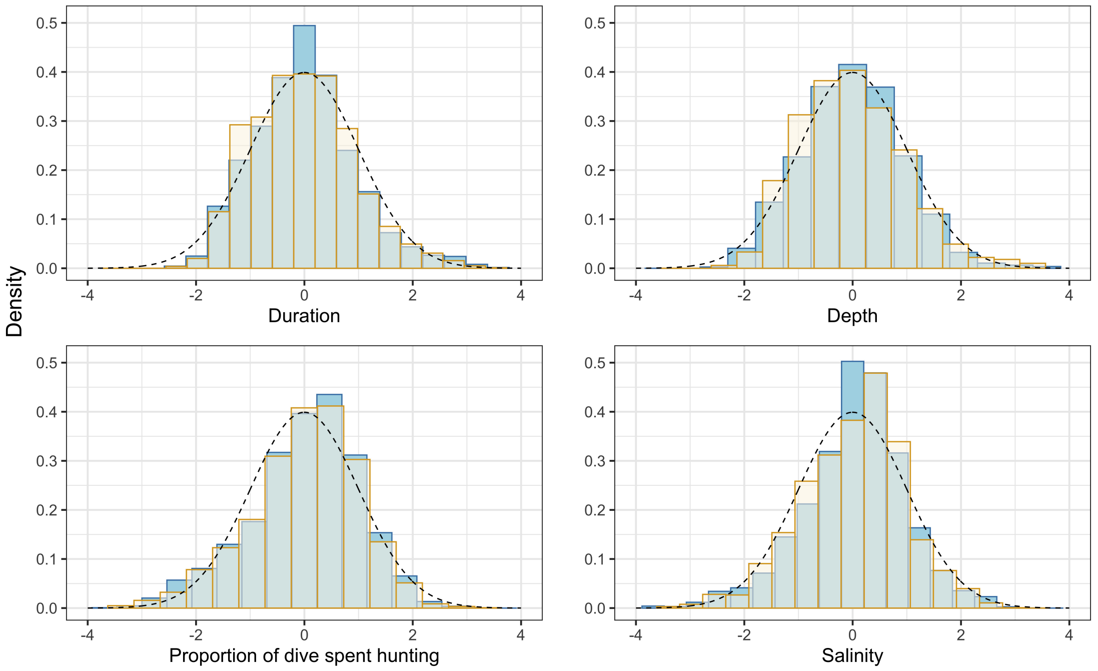


Figure S5.1. Histograms of the pseudo-residuals for each state-dependent variable in the model, excluding proportion of bathymetry which is an indicator variable. We show the pseudo-residuals for the female model in light blue, and pale yellow for the male model. The dashed black line shows the expected distribution of normally distributed residuals.

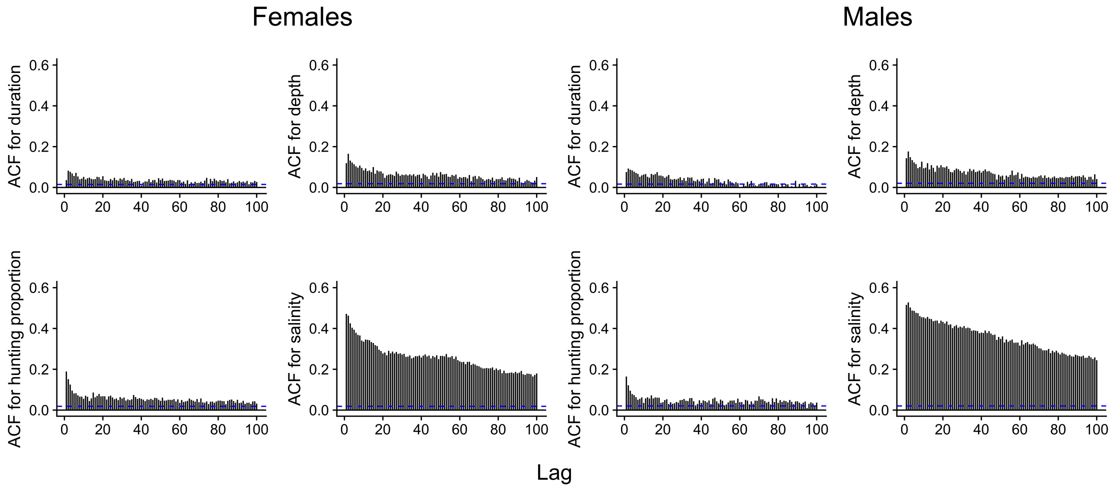


Figure S5.2. Autocorrelation plots of the pseudo-residuals for each state-dependent variable in the model, excluding proportion of bathymetry which is an indicator variable. The dashed blue line shows the 95% confidence interval around zero.

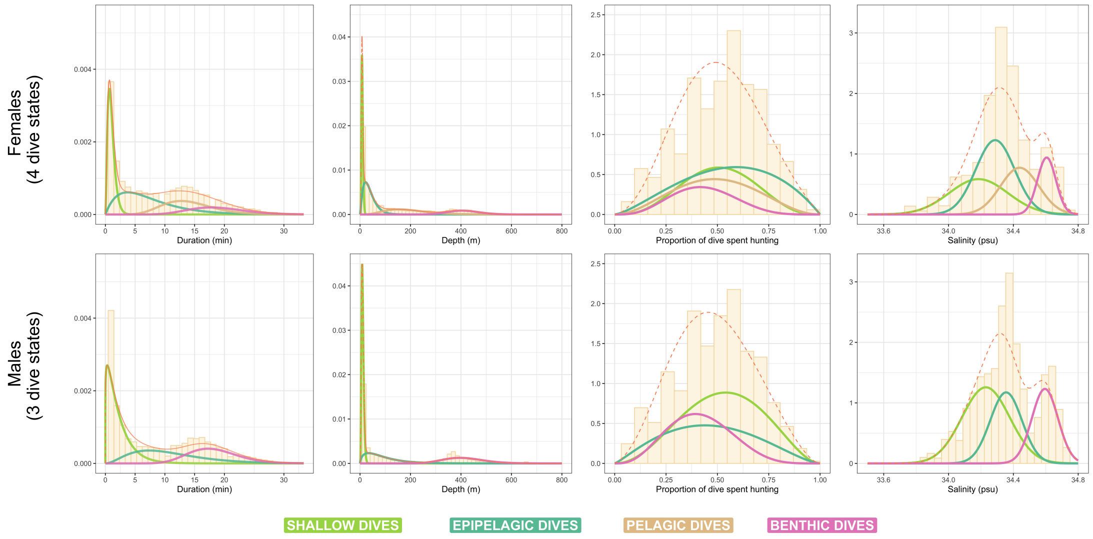


Figure S5.3. Histograms of the observed data for each state-dependent variable in the model, excluding proportion of bathymetry, which is an indicator variable. The component distributions are colour-coded by state and the marginal distribution is also shown in orange.

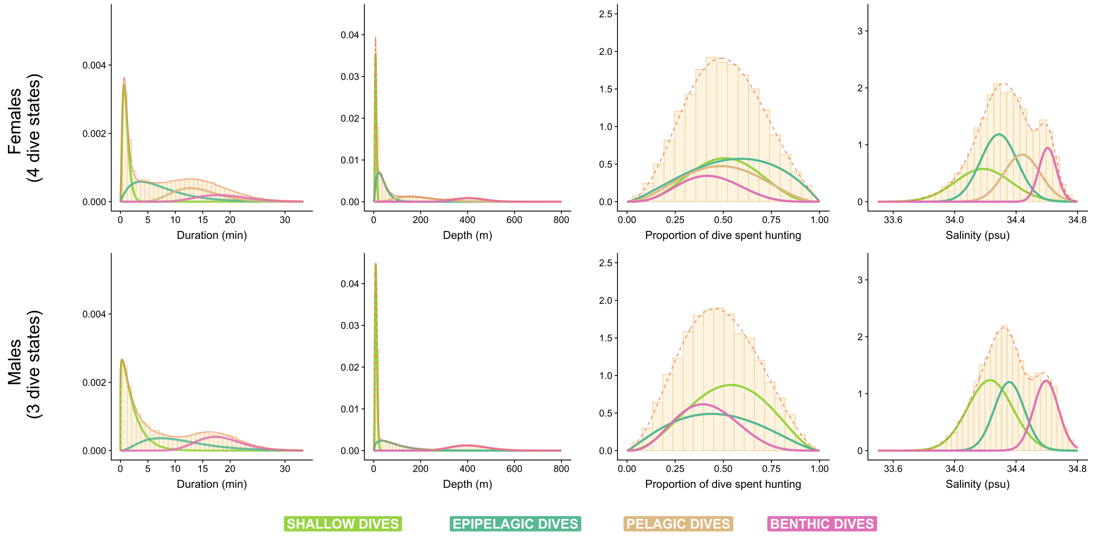


Figure S5.4. Histograms of the simulated data for each state-dependent variable in the model, excluding proportion of bathymetry, which is an indicator variable. The component distributions are colour-coded by state and the marginal distribution is also shown in orange.
